# Discrete cytokine signaling networks instruct distinct synovial pathotypes in inflammatory arthritis

**DOI:** 10.1101/2025.11.01.686006

**Authors:** Alicia Derrac Soria, David G. Hill, Stuart TO. Hughes, Reuben N. Scott, Ana Cardus Figueras, Sandra Dimonte, Daniela Costa, Ngoc-Nga Vinh, Federica Monaco, Robert H. Jenkins, Xiao Liu, Myles Lewis, Jason P. Twohig, Carol Guy, Aisling Morrin, Benjamin C. Cossins, Robert Andrews, Barbara Szomolay, Liliane Fossati-Jimack, Mari A. Nowell, Anwen S. Williams, Ernest H. Choy, Brendan J. Jenkins, Nigel M. Williams, Hua Yu, Marcin Kortylewski, Stephen Turner, Tony Tiganis, Costantino Pitzalis, Gareth W. Jones, Simon A. Jones

## Abstract

Patients with rheumatoid arthritis (RA) display distinct patterns of synovitis. To define the inflammatory mechanisms driving this heterogeneity, we analyzed the inflamed synovium of wild-type (WT), *Il6ra*^-/-^, and *Il27ra*^-/-^ mice with antigen-induced arthritis (AIA). Remarkably, each strain developed a joint pathology mirroring a major RA synovial pathotype: myeloid-rich (WT), fibroblast-rich/pauci-immune (*Il6ra*^-/-^), and lymphoid-rich (*Il27ra*^-/-^) synovitis. Histology confirmed minimal immune infiltration in *Il6ra*^-/-^ joints, while WT and *Il27ra*^-/-^ mice exhibited prominent immune involvement, including organized synovial lymphoid-like aggregates in *Il27ra*^-/-^ mice. Transcriptomic and epigenomic profiling revealed both shared and distinct regulatory programs among genotypes. *Il6ra*^-/-^ mice showed increased WNT, DKK, and AMPK signaling associated with fibroblast, chondrocyte, and osteoclast activation (e.g., *Adamts19*, *Dkk1*, *Ecm1*). Consistent with synovial ectopic lymphoid-like structures, *Il27ra*^-/-^ mice showed enrichment of lymphocyte activation (e.g., *Il17a*, *Il22*, *Bhlhe40*). WT mice exhibited hallmarks of MAP kinase activation. These molecular signatures parallel those of fibroblast-, lymphoid-, and myeloid-rich synovitis in RA. Defining a STAT1–STAT3 regulatory interplay influencing transcriptional decisions in WT and *Il27ra*^-/-^ mice, our findings offer insights into cytokine-driven disease heterogeneity. Together, these results establish a framework for mechanism-based classification of synovitis and introduce new mouse models to study the molecular drivers of synovial pathotypes and treatment response.

## Introduction

Biological and targeted drugs that block cytokine signals have greatly improved the treatment of immune-mediated inflammatory diseases (IMIDs) like RA (*1–3*). However, many patients do not respond well to certain treatments, which suggests that different cytokine networks affect the progression of arthritis (*2–4*). This limited effectiveness has driven advances in ultrasound-guided joint biopsies to study the biology of synovitis (*1–4*). These studies have shown that synovitis is highly variable and have led to clinical trials testing targeted therapies on specific synovial pathotypes (*5–8*). Yet, the inflammatory mechanisms behind this variability are still unclear. To address this, we have examined the gene expression programs that control synovitis by analyzing cytokine signaling in inflammatory joint disease.

Synovial biopsies from patients with RA show diverse cellular and molecular features of joint disease, allowing classification into fibroblast-rich (pauci-immune), myeloid-rich, and lymphoid-rich types of synovitis (*5, 9, 10*). It is currently unknown whether these pathotypes develop naturally over the disease course or arise from separate inflammatory mechanisms. Although cytokine-blocking drugs are established treatments for RA, few studies have examined how cytokines influence the development of specific synovial pathotypes in some patients but not others. For instance, little is known about how lymphocyte networks consisting of follicular dendritic and stromal cells form in lymphoid-rich synovitis, or how fibroblast-rich synovitis progresses without immune cell infiltration (*5, 11–16*).

In synovitis, pathological features such as cellular hyperplasia, osteoclastogenesis, cartilage erosion, and immune cell infiltration are driven by cytokines that often activate the Janus kinase (JAK)–Signal Transducer and Activator of Transcription (STAT) pathway (*17–20*). These discoveries have guided the development of drug interventions targeting this axis, including inhibitors of interleukin (IL)-6 (e.g., tocilizumab, sarilumab, sirukumab) and JAK proteins (e.g., tofacitinib, baricitinib) (*21–24*). Here, we aimed to elucidate how synovial heterogeneity is defined by cytokine-mediated signaling networks involving the JAK–STAT cascade. To this end, we compared the biological activities of IL-6 and IL-27, two cytokines that engage related receptor systems yet exert contrasting inflammatory outcomes.

Interleukin-6 and IL-27 activate the STAT1 and STAT3 transcription factors *via* receptor complexes that share gp130 as a common signaling β-subunit (*21, 25–30*). Despite this shared signaling architecture, the two cytokines frequently elicit opposing immunological outcomes, with IL-27 promoting STAT1-dependent programs that counterbalance STAT3-driven responses (*21, 25, 26, 31*). Pre-clinical models of arthritis underscore the functional relevance of this distinction, with *Il27ra*^-/-^ mice developing a severe pathology, whereas *Il6*^-/-^ mice display markedly attenuated disease (*19, 20, 32–38*). By characterizing the cellular and molecular signatures of joint inflammation in *Il6ra*^-/-^ mice and *Il27ra*^-/-^ mice with antigen-induced arthritis, we demonstrate that these genotypes recapitulate subsets of RA patients with fibroblast-rich and lymphoid-rich synovitis. Together, these findings reveal the mechanistic underpinnings of synovial heterogeneity and identify distinct cytokine pathways as potential targets for tailored therapeutic intervention.

## Materials and Methods

### Mice

Wild-type (WT) C57/Bl6 mice were sourced from Charles River UK. Mice deficient in *Il27ra* (B6N.129P2-Il27ra^tm1Mak^/J; The Jackson Laboratory) were generated by targeted disruption of the exon encoding the extracellular fibronectin type III domain of the IL-27R (also termed WSX-1) (*39*). Mice lacking *Il6ra* were established by ablating exons 4-6, which encode the IL-6 binding motifs (*40*).

### Antigen-induced arthritis

All studies of antigen-induced arthritis (AIA) were conducted under UK Home Office approvals (PPL-30/1820; PPL-30/2928; PB3E4EE13; PF34A3DC8; PE8BCF782) (*19, 20, 35, 38, 41*). Mice were challenged (s.c.) with methylated BSA (mBSA; 1 mg/ml emulsified in Complete Freund’s Adjuvant) and 160 ng *Bordetella pertussis* toxin (i.p.). Following an antigen boost with mBSA and CFA (s.c.), inflammatory arthritis was established by intra-articular (i.a.) administration of mBSA (10μl; 10 mg/ml) into the right knee joint. Synovial tissues were harvested on days 3 and 10 of AIA (*20*).

### Inhibition of synovial STAT3

The oligonucleotide design and synthesis of CpG-*Stat3*siRNA is described elsewhere (*42, 43*). Forward and reverse oligonucleotides (50 nmol) were dissolved in 250μl of DNase/RNase-free water at 37°C for 5-7 minutes before complementary hybridization by heating to 80°C for 1 minute and returning to 37°C for 1 hour. Mice were administered (i.a.) with CpG-*Stat3*siRNA (0.125 nmol/μl) at the point of arthritis induction in combination with mBSA (total volume of 10μl/joint). To test the specificity of this treatment, mice with AIA were also administered at the point of arthritis induction with either CpG alone or a non-hybridized, single-stranded version of CpG-*Stat3*siRNA.

### Histology

Knee joints were decalcified, formalin-fixed, and paraffin-embedded (HistoCore PEARL and Arcadia-H instruments; Leica Biosystems). Serial parasagittal sections (7μm) were stained in hematoxylin, fast green, and safranin-O. Synovitis was scored by three independent observers who evaluated sections for synovial infiltration (0–5), synovial exudate (0-3), synovial hyperplasia (0-3), and joint and cartilage erosion (0-3). An accumulative score of these parameters is presented as an Arthritis Index (*20, 35, 41*). All immunohistochemistry protocols were conducted as previously described.

### Transcriptomic analysis (RNA-seq)

Synovial RNA was isolated from tissue extracts using RNeasy Mini kits (QIAGEN). Libraries were generated from 2-4mg of mRNA following removal of cytoplasmic, mitochondrial, and ribosomal RNA (ribominus transcriptome isolation kit; Ambion). Libraries were prepared using the TrueSeq Stranded Total RNA library Prep kit (Illumina) or RNA-seq kit v2 (Life Technologies; 4475936) and sequenced on an Illumina HiSeq 4000 platform. Raw fastq files were processed using the nf-core/rnaseq (v3.17.0) pipeline with default parameters, and differentially expressed genes were analyzed with the DESeq2 package. The GRCm39 (release 112) reference was used for mapping. Genes that did not pass the log fold change or statistical significance threshold (adjusted *P*<0.05, log2FC >1 or log2FC <-1) were excluded from further analysis. Interpretation of gene function was performed using Ingenuity Pathway Analysis software (Qiagen), decoupleR, and the clusterProfiler functions enrichGO and enrichKEGG.

### Epigenetic analysis (ATAC-seq)

Genomic DNA was isolated from the synovium of mice with AIA. ATAC-seq was performed using 100,000 nuclei per sample as described in the manufacturer’s instruction protocol (Illumina, FC-121-1030). After amplification, library DNA was isolated (Qiagen MiniElute kit), size selected, and sequenced (Illumina HiSeq4000). All samples were quantified (Qubit; Invitrogen) before sequencing (Illumina HiSeq4000; 40-70M reads for each sample). Processing of raw reads was performed using nf-core/atacseq (v2.1.2). Two replicates per sample group were used to identify transcription factor DNA consensus peak sets. In all other cases, default parameters were applied. For mapping with BWA, the mouse GRCm39 reference genome was used. An occupational analysis was conducted with DiffBind to determine peak exclusivity. Before running DiffBind, variation in sample size was accounted for by ordering each sample by descending peak fold enrichment and retaining the first 20,000 peaks per sample. Consensus peak sets were produced using a minimum peak occurrence of 33% across the samples belonging to each condition. Pathway analysis of genes associated with peaks was performed using the annotatePeak function from ChIPseeker to align genes with sequencing peaks. The enrichGO function from clusterProfiler was used to identify pathways associated with these genes. Peaks occurring in the naïve WT control set were excluded, and remaining peaks were exported to bed format. Datasets were uploaded to MEME Suite and analyzed for motif enrichment using SEA (Simple Enrichment Analysis) using HOCMOCO Mouse (v11 CORE) motif database. Scanning of individual target motifs was conducted using FIMO (Find Individual Motif Occurrences). PFMs (Position Frequency Matrices) for each motif were obtained from the HOCOMOCO Mouse (v11 CORE) database. Default settings were applied to both SEA and FIMO.

### Quantitative PCR

Gene expression was quantified by TaqMan Gene Expression Assay (Applied Biosystems) using a barcoded MicroAmp Optical 384-well reaction plate (Applied Biosystems) (*20, 38*).

### Molecular Pathway Analysis and Modeling

Transcriptomic and epigenetic datasets were analyzed using Metacore integrated software suite (Thomson Reuters), Ingenuity Pathway Analysis (IPA; Ingenuity), and gene set enrichment analysis (GSEA). Genes were ranked by descending log2FC before enrichment was tested against STAT3 and STAT1 regulons defined by the DoRothEA database and retrieved through the decoupleR package. The GSEA function from clusterProfiler was used to test for enrichment. For machine learning analysis, genes selectively up-regulated (log2FC > 1, padj < 0.05) in the inflamed synovium of *Il6ra*^-/-^ and *Il27ra*^-/-^ mice were mapped to human orthologs induced in fibroid-rich and lymphoid-rich synovitis. The clusterProfiler package was then used to define gene modules curated according to defined biological processes. A total of 5 gene modules were identified from the gene lists established for fibroid-rich synovitis (*Il6ra*^-/-^) and lymphoid-rich synovitis (*Il27ra*^-/-^) mice. Each gene module was tested for predictive power using a generalized linear model with elastic net regularization and 5-fold cross-validation (glmnet package). Test-train splits in the data were determined using the caret package, with pROC used to visualize the ROC curves. K-means clustering (with 3 centers) overlayed PCA results with computationally derived clusters.

### Human samples and other repository datasets

Details of the clinical cohorts and metadata from studies of human synovial biopsies are previously described (*5, 7, 8*). Transcriptomic data from these studies (ArrayExpress:E-MTAB-6141) are publicly available (*5, 7*).

#### Visualization

Heatmaps were prepared using pheatmap and visualization packages in RColorBrewer and ggplot2. The gseaplot and dotplot functions were from the enrichplot package. Unless stated, heatmaps were clustered using the default complete linkage in pheatmap. Heatmaps of gene expression used Variance Stabilized Transformed counts, scaled in a row-wise manner, unless stated.

#### Statistics

Statistical comparisons of datasets used a two-tailed unpaired Student’s *t-*test. Unless stated, one-way ANOVA followed by Tukey’s comparison test was used for multiple comparisons. Analysis was conducted using GraphPad Prism 5 (GraphPad Software). Values showing \**P*< 0.05; \*\**P*< 0.01; \*\*\**P*< 0.001, \*\*\*\**P*< 0.0001 were considered significant.

## Results

### Mice lacking IL-6R or IL-27R develop synovial pathotypes resembling synovitis in rheumatoid arthritis

Patients with RA display joint pathologies that classify synovitis as fibroblast-rich, myeloid-rich, or lymphoid-rich (*5, 20*). These pathotypes range from low-inflammatory synovitis lacking infiltrating immune cells to pathologies with a diffuse or highly organized inflammatory infiltrate (Supplemental Figure 1). Previously, we identified that IL-27 limits the formation of synovial ectopic lymphoid-like structures (ELS) in AIA (*20*), raising the possibility that gp130 signaling through STAT1 and STAT3 may instruct pathological programs affecting synovitis heterogeneity.

To investigate the role of STAT1 and STAT3 in synovitis heterogeneity, we evaluated AIA in *Il6ra*^-/-^ and *Il27ra^-/-^* mice (**Figure 1**). These cytokine receptor cassettes signal through STAT1 and STAT3, with IL-27 showing a signaling bias for STAT1. When compared to WT mice with AIA, *Il6ra*^-/-^ and *Il27ra^-/-^* mice showed differences in antibody titers to mBSA challenge, joint swelling as a response to arthritis onset, and histological features of synovial pathology (**Figure 1A**; Supplemental Figure 1). Histology scores of arthritis (arthritis index) showed that *Il27ra*^-/-^ mice developed a more severe pathology than WT mice, with *Il6ra*^-/-^ mice displaying a more protected form of synovitis (**Figure 1A**). Scores for synovial infiltration, exudate, hyperplasia, and erosion showed that *Il6ra*^-/-^ mice with AIA developed synovitis with minimal evidence of immune cell involvement (**Figure 1A**). However, scores of synovial hyperplasia in *Il6ra*^-/-^ mice were equivalent to those of WT and *Il27ra*^-/-^ mice (**Figure 1A** & **1B**). To verify the stromal phenotype of synovitis in *Il6ra*^-/-^ mice with AIA, experiments were repeated in *Il6*^-/-^ mice using a relapsing/remitting model of AIA (Supplemental Figure 1). Recurrent activation of synovitis in *Il6*^-/-^ mice caused a progressive worsening of synovial hyperplasia that occurred without immune cell infiltration (Supplemental Figure 1). In contrast to *Il6ra*^-/-^ and *Il6*^-/-^ mice, WT and *Il27ra*^-/-^ mice with AIA showed leukocyte involvement, with synovitis in *Il27ra*^-/-^ mice characterized by synovial ELS (**Figure 1A** & **1B**; Supplemental Figure 1) as previously reported (*20*). These findings raise the possibility that gp130 cytokine receptor signaling through the Jak-STAT pathway shapes alternate patterns of synovitis in AIA.

**Figure 1.**
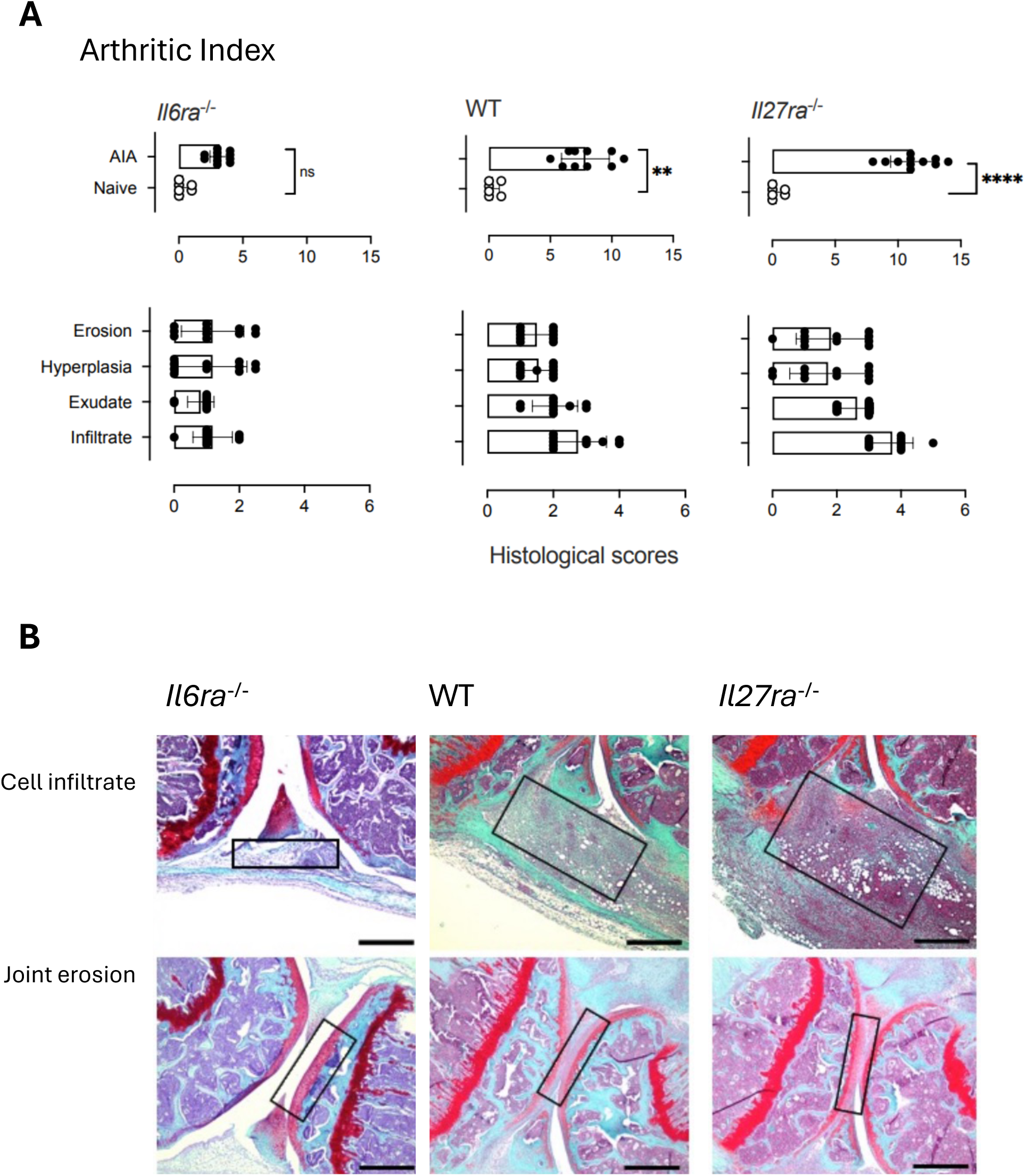
Mice with AIA display discernible features of disease heterogeneity. **(A)** WT, *Il6ra*^-/-^ and *Il27ra*^-/-^ mice were challenged with AIA (n=11 mice/group). Histology scores of disease activity are presented as an Arthritic Index (acquired at day 10 of AIA). Individual scores of synovial infiltrate, exudate, hyperplasia, and erosion are also shown. Values show scores from three independent assessors blinded to the study design (mean ± SEM, n=10-11 mice/group; \*\**P*<0.01; \*\*\*\**P*<0.001). **(B)** Representative hematoxylin and safranin-O staining of knee joints taken on day 10 of AIA (scale bar, 100μm). Boxes show cellular infiltration (top panels) and erosion (bottom panels).

### Mouse models of AIA display common and unique patterns of synovial gene regulation

Next, we conducted RNA-seq of synovial tissues harvested from the synovium of control mice before disease and at days 3 and 10 of AIA (**Figure 2**, Supplemental Figure 2). Bioinformatic analysis of transcriptomic data from WT, *Il6ra*^-/-^, and *Il27ra*^-/-^ mice with AIA revealed gene changes following the onset of synovitis (Supplemental Figure 2). Consistent with our histological determinations, GSEA identified pathways associated with activated fibroblasts, monocytic cells, and lymphocytes in the datasets from WT, *Il6ra*^-/-^, and *Il27ra*^-/-^ mice, respectively (**Figure 2A**). However, we also observed gene signatures common to all three mouse strains with AIA.

**Figure 2.**
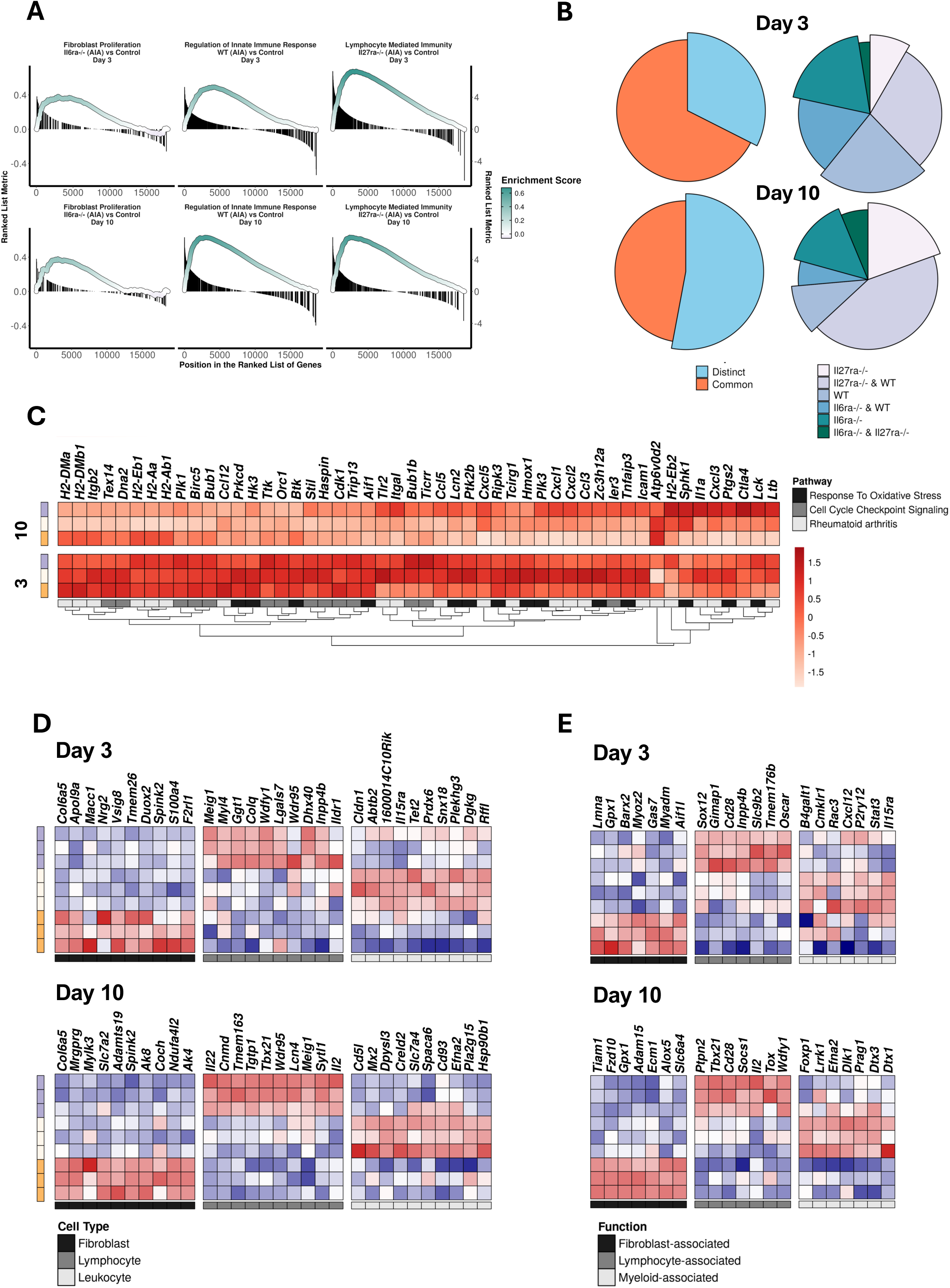
WT, *Il6ra*^-/-^ and *Il27ra*^-/-^ mice with synovitis display common and unique transcriptional signatures. Bulk RNA-seq was performed on synovial tissues extracted from WT, *Il6ra*^-/-^, and *Il27ra*^-/-^ mice at baseline and days 3 and 10 of AIA. For each condition, transcriptomic data were averaged from 3 independent mice. See Supplemental Figure 1 for genes differentially regulated in response to AIA displayed as a heatmap (Log2FC>1; adjPvalue<0.05). Complete linkage was applied to create a hierarchical clustering of genes. Gene expression was ranked according to Log2FC and scaled according to z-score. **(A)** Differentially regulated gene sets from *Il6ra*^-/-^, WT, and *Il27ra*^-/-^ mice with AIA were expressed relative to naïve non-challenged mice and analyzed by Gene Set Enrichment Analysis (GSEA). Results for days 3 and 10 of AIA are shown, identifying pathways integral to fibroblast, myeloid, and lymphocyte-driven pathologies. **(B)** Pie charts showing the proportion of arthritis-inducible genes common (orange/brown) and unique (purple/blue) to each mouse strain. A separate pie chart focuses on uniquely regulated genes, showing the proportion of genes expressed by an individual mouse strain and those shared with another strain at days 3 and 10 of AIA. **(C)** Examples of genes common to all three mouse strains are displayed as heatmaps. Processes linked to their biological function were identified by pathway analysis (clusterProfiler). **(D)(E)** Heat maps show examples of genes defining the features of synovitis in WT, *Il6ra*^-/-^, and *Il27ra*^-/-^ mice with AIA at day 3 and 10 of AIA (Log2FC>1; adjPvalue<0.05). Phenotypic markers identifying fibroblasts, myeloid cells, and lymphoid cells **(D)** and pathways associated with their activation **(E)** are shown.

The proportion of transcripts shared between the strains was more apparent at the early stage of AIA (day 3), but a sizeable number were also seen at day 10 (**Figure 2B**). These include genes linked to innate sensing (e.g., *Il1a*, *Tlr2*, *Tlr4*), cell adhesion (e.g., *Icam1*), DNA repair (e.g., *Cdk1*, *Dna2*, *Tex14*), oxidative stress (e.g., *Hk3*, *Hmox1*, *Ripk3*), and inflammatory indices including cytokines (e.g., *Il15*, *Il18*, *Il18bp*), chemokines (e.g., *Ccl3*, *Ccl5*, *Cxcl2*), and regulators of cellular proliferation (**Figure 2C**; Supplemental Figure 2). Despite these similarities, synovial tissues from WT, *Il6ra*^-/-^, and *Il27ra*^-/-^ mice with AIA showed differences in synovial gene expression (**Figure 2B**). These were more evident on day 10 of AIA. Genes enriched in *Il6ra*^-/-^ mice included those associated with stromal cell activation, including those involved in energy metabolism and tissue homeostasis (e.g., *Ak4*, *Ak8*, *Slc7A2*, *Spink2*, *S100a4*) (**Figure 2D**). In contrast, synovitis in WT and *Il27ra*^-/-^ mice with AIA revealed evidence of genes controlling leukocyte functions. These include markers governing the maintenance, differentiation, or survival of monocytic cells in WT mice (e.g., *Cd93*, *Il15ra*, *Prdx6*, *Snx18*), and lymphoid effector functions (e.g., *Il2*, *Il22*, *Tbx21*) in *Il27ra*^-/-^ mice with AIA (**Figure 2D**). Consistent with these results, we identified CXC- and CC-chemokines and their receptors governing defined patterns of leukocyte infiltration (Supplemental Figure 2). For example, synovial *Cxcl12*, *Cxcl13*, and *Ccl19,* involved in ELS formation, were upregulated in synovial tissues from *Il27ra*^-/-^ mice (Supplemental Figure 2). Thus, WT, *Il6ra*^-/-^, and *Il27ra*^-/-^ mice display molecular signatures of cell types controlling synovitis heterogeneity.

To understand how genes uniquely expressed in WT, *Il6ra*^-/-^, and *Il27ra*^-/-^ mice with AIA contribute to synovitis, we performed molecular pathway analysis to identify signatures of stromal and leukocyte activation (**Figure 2E**). Pathways controlling fibroblasts, chondrocytes, and osteoclasts were enriched in *Il6ra*^-/-^ mice with AIA. These included genes contributing to synovial hyperplasia (e.g., *Fzd10*, *Gpx1*), or bone (e.g., *Adam15*, *Barx2*) and cartilage (e.g., *Ecm1*, *Lmna*, *Tiam1*) erosion (**Figure 2E**). By contrast, WT and *Il27ra*^-/-^ mice with AIA were more closely associated with pathways affecting leukocyte recruitment, activation, or retention. WT mice with AIA showed a bias towards pathways regulating innate signaling or myeloid cell differentiation and maturation (e.g., *Dtx1*, *Dtx3*, *Dlk1*, *Efna2*, *Foxp1*, *Prag1*) (**Figure 2E**), whereas *Il27ra*^-/-^ mice with AIA displayed heightened lymphocyte activation (e.g., *Cd28*, *Il2*, *Ptpn2*, *Tox*) (**Figure 2E**). These findings support the histology in Figure 1 and open possibilities to define the transcriptional programs behind synovitis heterogeneity.

### Distinct signaling pathways instruct the development of synovial pathotypes

To investigate the regulatory pathways triggered by AIA in WT, *Il6ra*^-/-^, and *Il27ra*^-/-^ mice, we used decoupleR. Analysis revealed the dynamic and contrasting nature of signaling events activated in each mouse strain (**Figure 3**). These included cytokines (e.g., TNFα, TRAIL/TNFSF10), growth factors (e.g., EGF, TGFβ, VEGF), and connections to NF-κB, Jak-STAT, MAP kinase, HIF, WNT, and PI3-kinase signaling pathways (**Figure 3A**, Supplemental Figure 2). Key differences in the synovial expression of signaling intermediates controlling these cascades were observed at days 3 and 10 of AIA (**Figure 3B**). Compared to WT or *Il6ra*^-/-^ mice with AIA, *Il27ra*^-/-^ mice showed a more restricted synovial expression of signaling molecules at day 3 of synovitis (**Figure 3B**). This profile was reversed by day 10 of AIA, with synovial tissues from *Il27ra*^-/-^ mice displaying a more robust expression of genes. These included an increased expression of feedback inhibitors (e.g., *Nfkbia*, *Nfkbib*, *Socs1*, *Socs3*), suggesting that these pathways were active in synovitis (Supplemental Figure 2). In contrast, *Il6ra*^-/-^ mice showed enrichment of signaling intermediates associated with fibrosis, hyperproliferation, and bone and cartilage turnover (e.g., *Wnt2*, *Wnt4*, *Dkk1*, *Dkk2*, *Irf4*, *Irf6*) (**Figure 3B** & Supplemental Figure 2). Upstream regulator analysis in IPA verified these findings (Supplemental Figure 2; see Supplemental Tables 1-3 for statistical summaries). Examples of cytokines, kinases, and transcription factors are presented for *Il6ra*^-/-^ and *Il27ra*^-/-^ mice, with ranked enrichment plots depicting activation z-scores relative to WT mice at days 3 and 10 of AIA (**Figure 3B**). Consistent with a lymphoid-driven synovitis, we identified pathways that regulate lymphocyte activities in *Il27ra*^-/-^ mice. These included lymphokines (e.g., *Il2*, *Il21*, *Ifng*) and markers of Th17 differentiation (e.g., *Baft*, *Bhlhe40*, *Fosl1*, *Il17a*, *Il23*, *Rorc*) or lymphocyte activation (e.g., *Cd40*, *Fos*, *Icos*, *Junb*) (**Figure 3B**). In contrast, *Il6ra*^-/-^ mice with AIA showed enrichment of cytokines (e.g., *Il4*, *Il11*, *Il13*), and Jak proteins (e.g., *Tyk2*), with functions affecting cell renewal, bone turnover, tissue homeostasis, and neovascularization (**Figure 3B**; Supplemental Figure 2). These include regulators (e.g., *Adam12*, *Adam17*) and downstream effectors of DKK, GATA6, NOTCH/Wnt, PPARγ, or TGFβ/BMP7 signaling (Supplemental Figure 2). To illustrate the impact of IL-4/IL-13 and IL-2/IL-17 on synovitis in *Il6ra*^-/-^ and *Il27ra*^-/-^ mice, we used IPA to generate heatmaps of gene signatures reflecting the activity of these cytokines in AIA (**Figure 3C**).

**Figure 3.**
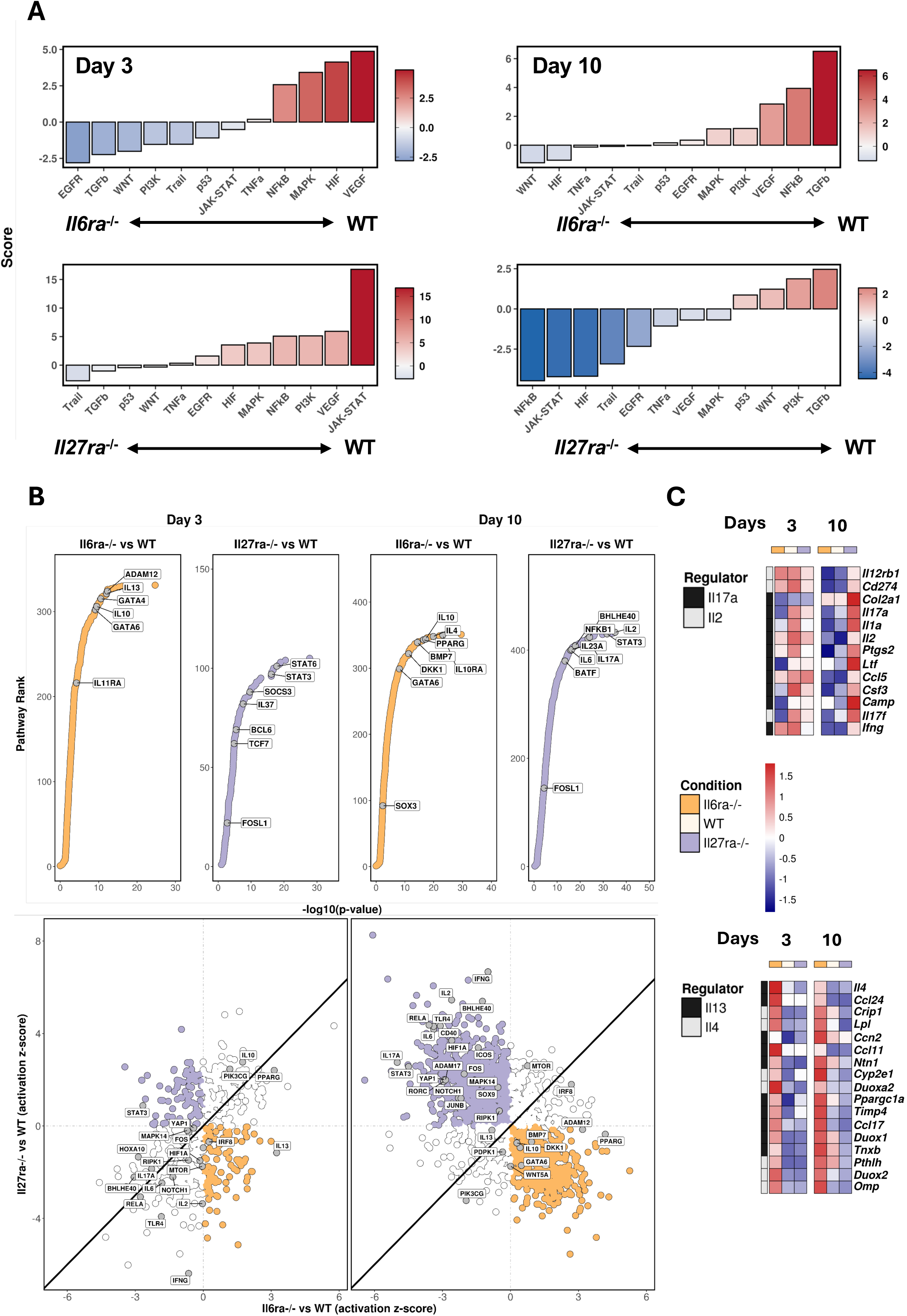
Regulatory signatures of synovitis in mice with AIA. **(A)** Bar plots depict the activation of signaling pathways identified using decoupleR in synovial tissues from WT, *Il6ra*^-/-^, and *Il27ra*^-/-^ mice with AIA. Results are shown for days 3 (left-hand panels) and 10 (right-hand panels) of AIA. A positive score indicates pathways with an increased enrichment in WT mice. **(B)** Ranked enrichment plots (top panels) show upstream regulators identified for *Il6ra*^-/-^ and *Il27ra*^-/-^ mice with AIA. The correlation plots (below panels) show upstream regulators comparisons between synovitis in *Il6ra*^-/-^ mice with AIA (orange) and *Il27ra*^-/-^ mice with AIA (purple). In both cases, data are expressed relative to WT mice with AIA. **(C)** Heatmaps of gene signatures of IL-2, IL-4, IL-13, and IL-17 activity identified by upstream regulator analysis in IPA (Log2FC>1; adjPvalue<0.05).

To verify the upstream regulator analysis, ATAC-seq of the inflamed synovium was performed to map changes in chromatin accessibility in response to AIA (**Figure 4**). Next-generation sequencing peaks were mapped to genomic loci and aligned with corresponding RNA-seq data (Supplemental Figure 3). There was a close association between gene annotations identified by ATAC-seq and RNA-seq, with datasets showing a 75-90% overlap (**Figure 4A**). Molecular pathway analysis of these shared signatures identified biological processes consistent with those listed in Figures 2 and 3 (**Figure 4B**, data not shown). Thus, ATAC-seq provides an opportunity to understand the transcriptional mechanisms underlying synovitis in WT, *Il6ra*^-/-^, and *Il27ra*^-/-^ mice with AIA. Simple Enrichment Analysis (SEA) of DNA motifs underlying next-generation sequencing peaks identified consensus sequences for transcription factors shared among all three mouse strains. These included zinc finger transcriptional regulators (e.g., CTCF, KLF, ZBTB, SP1 family), components of the AP1 system (e.g., FOS, FOSL1, FOSL2, FOSB, JUNB), basic region/leucine zipper superfamily members (e.g., ATF, CREB), and interferon response factors regulating myeloid and stromal cell activities (e.g., IRF3, IRF8) (**Figure 4C**; Supplemental Figure 3).

**Figure 4.**
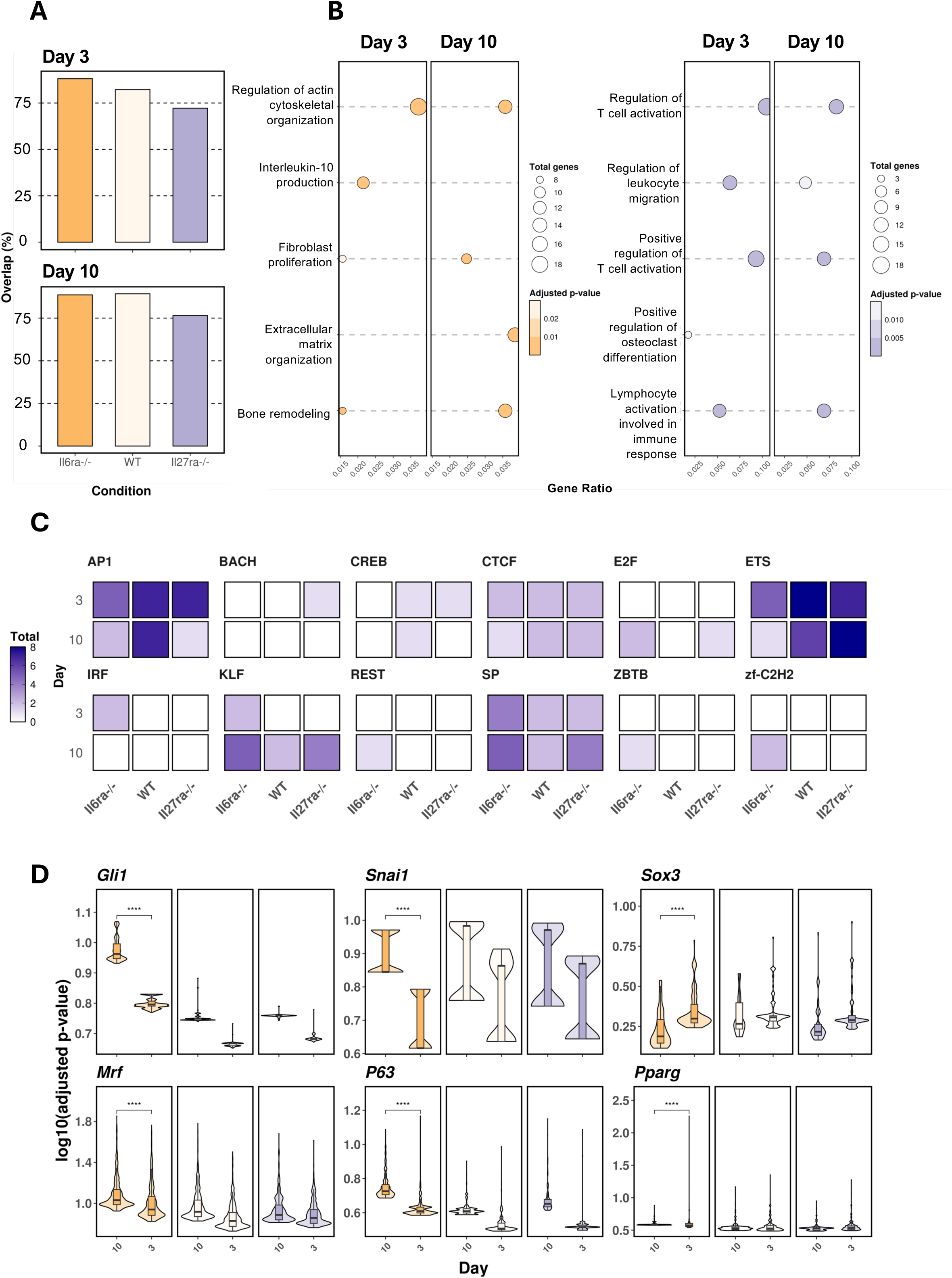
Epigenetic analysis of synovial inflammation from mice with AIA. **(A)** Genomic DNA was extracted from the inflamed synovium of WT, *Il6ra*^-/-^, and *Il27ra*^-/-^ mice with AIA for analysis by ATAC-seq. Consensus (≥2) ATAC-seq peaks were mapped (ChIPseeker) against annotated RNA-seq datasets to align changes in chromatin accessibility with alterations in gene expression. The percentage overlap between ATAC-seq and RNA-seq datasets is shown. **(B)** Molecular pathway analysis of transcripts sharing changes in chromatin architecture for *Il6ra*^-/-^ and *Il27ra*^-/-^ mice at day 3 and day 10 of AIA. **(C)** Putative transcription factor motifs associated with sequencing peaks from ATAC-seq datasets were identified using motif enrichment analysis in MEME-Suite (see Supplemental Figure 3). Heatmaps show the presence or absence of these sequences in ATAC-seq data from WT, *Il6ra*^-/-^, and *Il27ra*^-/-^ mice with AIA. **(D)** Violin plots showing access to other transcription factor motifs in ATAC-seq datasets. These were selected based on the upstream regulator analysis (IPA) of RNA-seq datasets from WT, *Il6ra*^-/-^, and *Il27ra*^-/-^ mice with AIA.

Based on the identification of DKK, GATA6, NOTCH/Wnt, PPARγ, and TGFβ/BMP7 signaling signatures in synovial tissues from *Il6ra*^-/-^ mice (Supplemental Figure 2), we tested whether transcription factors linked to these pathways were present in ATAC-seq datasets from *Il6ra*^-/-^ mice with AIA. Consensus motifs for TGFβ (GLI1, SNAI3), WNT (MYOD, MYOG, SOX3), NOTCH (P63), and PPARγ signaling were identified as open in all three mouse strains with AIA (**Figure 4D**). However, these loci were statistically more enriched in *Il6ra*^-/-^ mice with AIA, upholding the earlier computational predictions (**Figure 3**, **Figure 4D**, Supplemental Table 1-3). These predictions offer new insights into the molecular hallmarks affecting the onset and temporal regulation of synovial disease heterogeneity.

### Mice with AIA share molecular hallmarks of synovial pathotypes in rheumatoid arthritis

To explore connections with human pathology, we next compared our mouse findings to RNA-seq from joint biopsies of rheumatoid arthritis patients with known synovial pathotypes (**Figure 5**, Supplemental Figure 4). Pathway enrichment analysis identified that fibroblast-rich synovitis shares biological processes with those observed in *Il6ra*^-/-^ mice with AIA, whereas lymphoid-rich synovitis revealed pathways consistent with *Il27ra*^-/-^ mice with AIA (**Figure 5A**). To substantiate these findings, we analyzed mouse fibroblast and lymphoid gene sets presented in Figure 2 for their expression in human synovitis (**Figure 5B**). We observed a close relationship between fibroblast genes enriched in *Il6ra*^-/-^ mice with AIA and fibroblast-rich synovitis, as well as lymphocyte genes in *Il27ra*^-/-^ mice with AIA and lymphoid-rich synovitis (**Figure 5B**). Circos plots illustrate the association of all differentially regulated gene sets from WT, *Il6ra*^-/-^, and *Il27ra*^-/-^ mice with AIA to those of myeloid-rich, fibroblast-rich, or lymphoid-rich synovitis, respectively (**Figure 5C**). A heatmap displaying the expression of genes in human synovial biopsies characterized as myeloid-rich, lymphoid-rich, and fibroblast-rich synovitis highlights commonalities between mouse and human datasets (**Figure 5D**). In a separate approach, we applied machine learning methods to verify the relationship between mouse and human datasets (**Figure 6**). Extracting results from this analysis, we identified 91 transcripts as selectively up- regulated in fibroblast-rich synovitis and *Il6ra*^-/-^ with AIA or lymphoid-rich synovitis and *Il27ra*^-/-^ mice with AIA (**Figure 6A**). Principal component analysis of expression data from each human synovial biopsy is shown, identifying the histological classification of individual biopsies as fibroblast-rich, myeloid-rich, or lymphoid-rich (**Figure 6B**). K-means clustering was used to define three groupings sharing a statistical relationship, and the proportion of each synovial pathotype residing within each cluster is shown (**Figure 6B**). These data illustrate the potential fluid nature of myeloid-rich synovitis, with biopsies classified as this pathotype sharing features aligned to either fibroblast-rich or lymphoid-rich. Aligned to this approach, we used molecular pathway analysis to reveal genes sharing common biological processes. Gene predictions from this analysis were used to define gene modules from the top 5 hits for *Il6ra*^-/-^ with AIA and *Il27ra*^-/-^ mice with AIA (**Figure 6C**). Each gene module was tested against transcriptomic data from human synovial biopsies for predictive power using a generalized linear model with elastic net regularisation and 5-fold cross-validation (**Figure 6C**). This machine learning approach confirmed the similarity between the mouse and human datasets, with the ROC curves showing the power of each gene module to predict lymphoid-rich or fibroblast-rich synovitis (**Figure 6C**).

**Figure 5.**
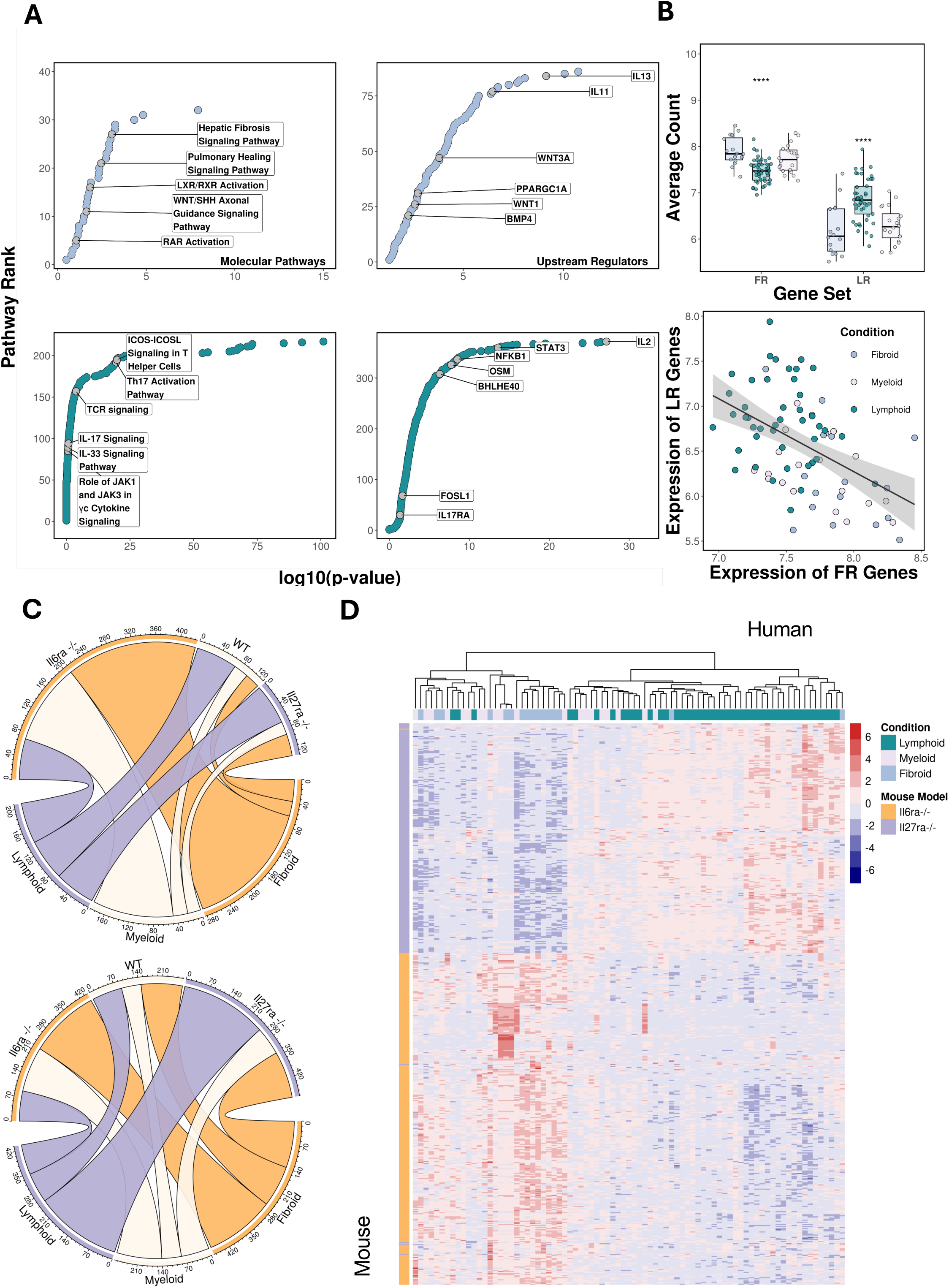
Mouse and human pathotypes share common transcriptional profiles. **(A)** Bioinformatic analysis of RNA-seq from synovial biopsies of patients with rheumatoid arthritis. Ranked enrichment plots show molecular pathways and upstream regulators ranked according to statistical significance. Data is shown for fibroblast-rich and lymphoid-rich synovitis relative to changes in myeloid-rich synovitis. **(B)** Murine gene lists of phenotypic fibroblast or lymphocyte markers (derived from Figure 2) were analyzed in RNA-seq data of lymphoid-rich, myeloid-rich, and fibroblast-rich synovitis. Box plots show the expression of individual transcripts (left) and their distribution across samples as a correlation (right). **(C)** Circos plots showing shared relationships between RNA-seq from WT, *Il6ra*^-/-^, and *Il27ra*^-/-^ mice with AIA and human myeloid-rich, fibroblast-rich, and lymphoid-rich synovitis. **(D)** Hierarchical clustering of genes extracted from the circos plots.

**Figure 6.**
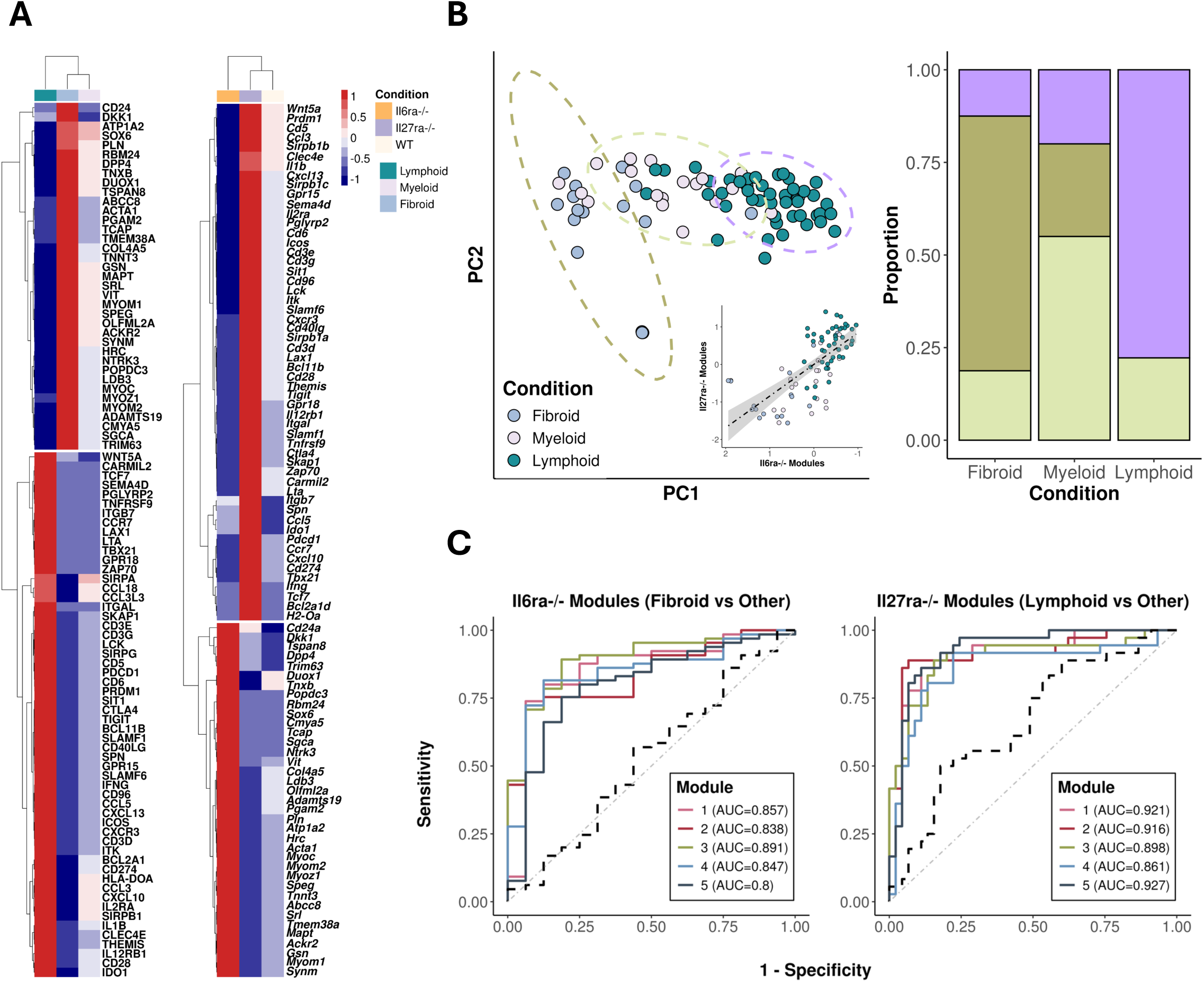
Prediction classifiers confirm the relationship between mouse and human transcriptomics. **(A)** Heatmaps show the relative expression of genes selectively upregulated in *Il6ra*^-/-^ and *Il27ra*^-/-^ mice with AIA (day 10). Corresponding gene expression data from fibroid-rich and lymphoid-rich synovitis are also shown. **(B)** PCA plot clustering of human data based on the expression of overlap genes identified in mice with AIA. K-means clustering into 3 centers was used to overlay PCA with clusters. Correlation between *Il6ra*^-/-^ and *Il27ra*^-/-^ gene modules is shown based on their relative expression in human transcriptomic data from synovial biopsies. In both cases, dots are color-coded to define each synovial pathotype. The stacked bar graph shows the proportion of synovial biopsies classified as fibroblast-rich, myeloid-rich, or lymphoid-rich synovitis falling into each k-means cluster. **(C)** Molecular pathway analysis of the gene lists shown in Panel A was used to assign their function to biological processes, creating defined gene modules for testing as disease classifiers. Receiver Operating Characteristic (ROC) curves of the performance of the top 5 modules for each condition, and Area Under the Curve (AUC) statistics, are shown. As a control, we constructed a module from genes displaying minimal variation in expression between all three pathotypes (presented as a dotted line; AUC 0.53 and 0.67 for fibroblast-rich and lymphoid-rich, respectively).

Building on this approach, we interrogated human and mouse datasets to understand the biological processes behind each synovial pathotype. Pathway analysis of gene sets identified links to fibrosis, the turnover of extracellular matrix, proteolytic shedding events, and the control of osteoclasts, osteoblasts, and chondrocytes (**Figure 7A**). In contrast, *Il27ra*^-/-^ mice with AIA and lymphoid-rich synovitis shared pathways regulating leukocyte migration and lymphocyte effector functions (**Figure 7A**). We further identified similar signaling mechanisms controlling human synovial pathotypes as those defined by upstream regulator analysis in AIA (**Figure 7B**). Correlation plots reveal the regulatory components common to fibroblast-rich synovitis and *Il6ra*^-/-^ mice or between lymphoid-rich synovitis and *Il27ra*^-/-^ mice with AIA (**Figure 7B**). These include the enrichment of WNT, DKK, TR/RXR, and AMPK signaling pathways, along with signatures of IL-11, IL-13, and Tyk2 activity in *Il6ra*^-/-^ mice with AIA and fibroblast-rich synovitis (**Figure 7B**). Signaling pathways involving NF-κB, C/EBPβ, STAT3, and cytokines such as TNFα, oncostatin-M, IL-1β, and IL-17 were all suppressed in *Il6ra*^-/-^ mice with AIA and fibroblast-rich synovitis (**Figure 7B**). Both *Il27ra*^-/-^ mice with AIA and lymphoid-rich synovitis showed positive correlations with lymphokines (e.g., IL-2, IL-6, IL-7, IL-12, IL-21, IL-23) and effector cytokines (e.g., IL-17A, IL-17F, IL-22). These signatures were significantly enriched in synovial tissues from day 10 of synovitis in *Il27ra*^-/-^ mice with AIA (**Figure 7C**, Supplemental Figure 4). To highlight this relationship, we compared the regulatory involvement of IL-2 and IL-13 in human and mouse datasets using findings presented in Figure 3C. Statistical associations showed a link between signatures of IL-2 activity in *Il27ra*^-/-^ mice with AIA and those identified in lymphoid-rich synovitis (**Figure 7D**). We observed a similar trend between markers of IL-13 involvement in *Il6ra*^-/-^ mice with AIA and fibroblast-rich synovitis (**Figure 7D**). Thus, the development of AIA in *Il6ra*^-/-^ and *Il27ra*^-/-^ mice exhibits similarities to the hallmarks of fibroblast-rich and lymphoid-rich synovitis in humans.

**Figure 7.**
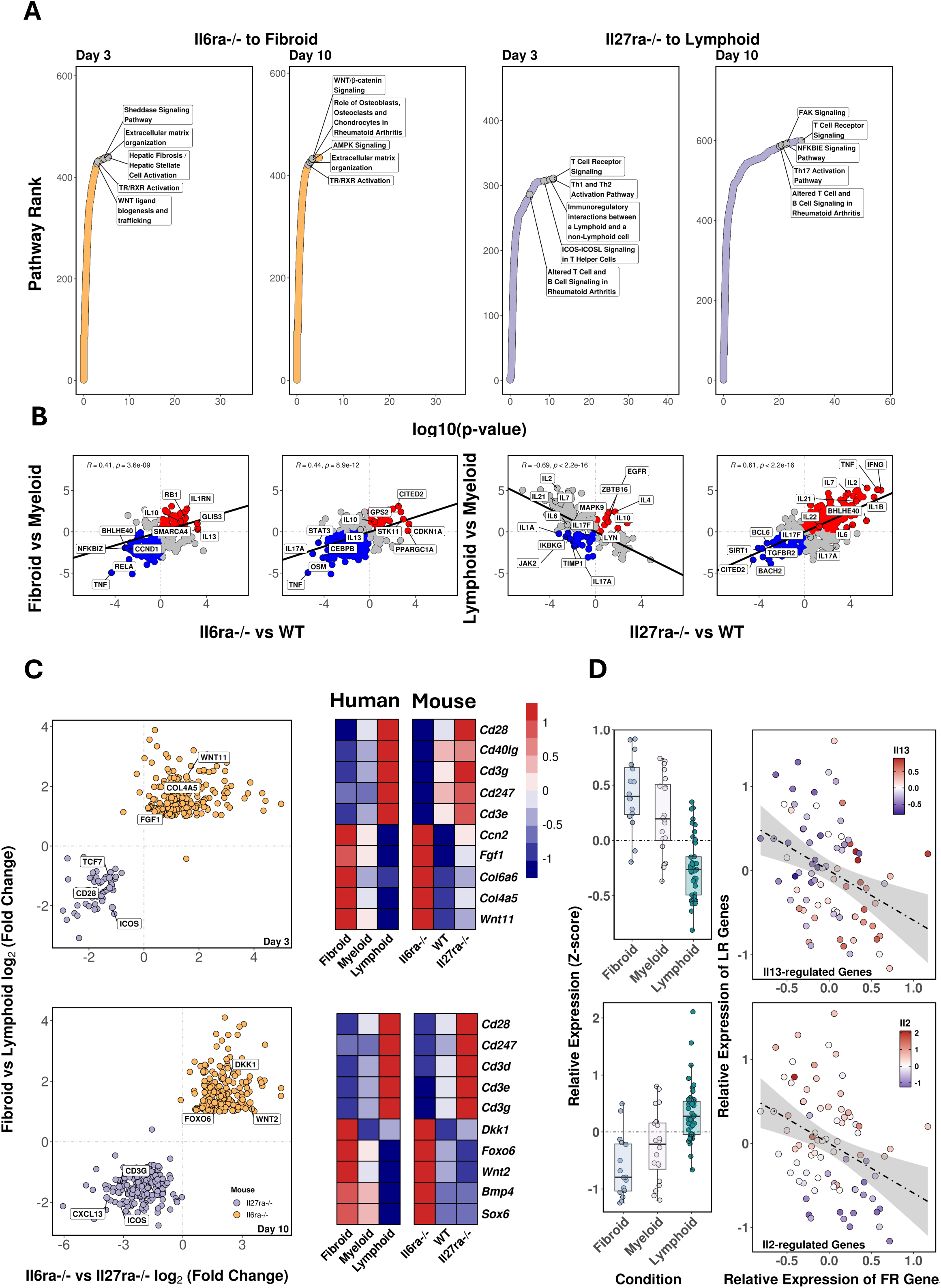
Mouse and human pathotypes display common upstream signaling programs. **(A)** Molecular pathway analysis of gene transcripts shared between *Il6ra*^-/-^ mice with AIA and fibroblast-rich synovitis, and *Il27ra*^-/-^ mice with AIA and human lymphoid-rich synovitis. **(B)** Results of upstream regulator analysis are used to produce correlation plots. *Il6ra*^-/-^ and *Il27ra*^-/-^ mice are compared to WT, whilst fibroid- and lymphoid-rich synovitis are compared to myeloid-rich synovitis. **(C)** Expression of shared gene signatures in molecular pathways from Panel A. Plots show differentially regulated transcripts between *Il6ra*^-/-^ and *Il27ra*^-/-^ mice with AIA at days 3 and 10 AIA. Heatmaps based on relative gene expression provide illustrative examples of genes shared between mice with AIA and human synovitis (see Supplemental Figure 6 for others). **(D)** Activation z-scores of gene signatures for IL-13 and IL-2 associated with phenotypic markers of fibroblasts and lymphocytes (as in Figure 2). Box plots show the relative expression of IL-2 and IL-13-regulated genes.

### Synovial interplay between STAT1 and STAT3 instructs synovitis heterogeneity

To test whether mice with AIA could help dissect the pathological processes driving synovitis heterogeneity, we focused on the transcriptional mechanisms governed by IL-6 and IL-27. We first applied the FIMO tool within MEME Suite to identify chromatin regions with accessible consensus sequences (TTCNGAA) for STAT transcription factor binding (**Figure 8A**). FIMO analysis revealed that promoters containing these DNA motifs were open (or poised) for transcription at day 3 of AIA. Access to these sites was maintained as synovitis developed in WT and *Il27ra*^-/-^ mice with AIA (**Figure 8A**). This was not seen in *Il6ra*^-/-^ mice with AIA (**Figure 8A**). Instead, chromatin regions containing consensus sequences for STAT transcription factor binding were closed by day 10 of AIA in *Il6ra*^-/-^ mice (**Figure 8A**). These data suggest that IL-6 signaling regulates chromatin accessibility. In support, we observed an enrichment of consensus motifs for the RE1 silencing transcription factor (REST) in ATAC-seq datasets from *Il6ra*^-/-^ mice with AIA (**Figure 4C**).

**Figure 8.**
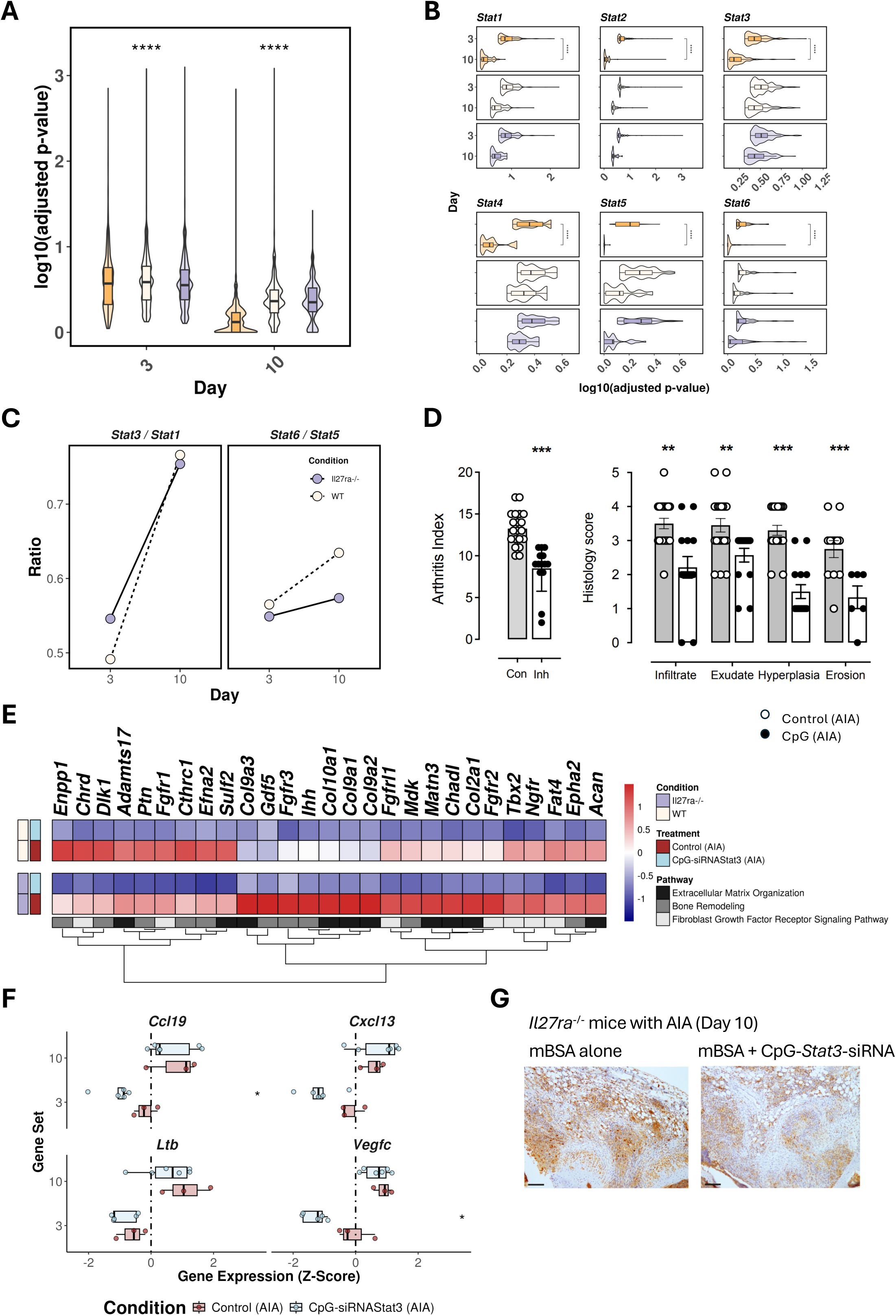
Profiling of STAT transcription factor involvement in mice with AIA. **(A)** Violin plots reported the incidence of STAT transcription factor consensus sequences in ATAC-seq datasets at days 3 and 10 of AIA. Statistics are based on the Kruskal-Wallis test. Summary statistics are shown in Supplemental Table 1. **(B)** Violin plots show the frequency of consensus sequences for the STAT transcription factor family in ATAC-seq datasets. Sites were identified using motif enrichment analysis in MEME-Suite, with data shown for WT, *Il6ra*^-/-^, and *Il27ra*^-/-^ mice at days 3 and 10 of AIA. **(C)** Graphs show the STAT3:STAT1 and STAT6:STAT5 ratios at day 3 and 10 of AIA based on the frequency of mapped consensus sequences from Panel A. Summary statistics are shown in Supplemental Tables 2 & 3. **(D)** Mice were primed for AIA and, on disease induction, administered (i.a.) with mBSA alone or with CpG-*Stat3*-siRNA (0.125 nmol/μl). The Arthritis Index and scores of synovial infiltration, exudate, hyperplasia, and joint damage are shown (mean ± SEM; n=12-14 per group). Mice treated with CpG alone or a non-hybridized, single-stranded variant of CpG-*Stat3*-siRNA were used as controls (see Supplemental Figure 5). **(E)** RNA-seq analysis of synovial tissues for WT and *Il27ra*^-/-^ mice with AIA. Analysis showing representative gene changes at day 10 of AIA for mice treated with mBSA alone or mBSA and CpG-*Stat3*-siRNA (Log2FC>1; adjPvalue<0.05). **(F)** Relative expression of genes involved in the organization and maintenance of ectopic lymphoid-like structures in *Il27ra*^-/-^ mice with AIA. *Ccl19*, *Cxcl13*, *Ltb*, and *Vegfc* expression is shown for mice treated with mBSA alone or with CpG-*Stat3*-siRNA. **(G)** Representative immunohistochemistry of CD3 in sections taken at day 10 of AIA from *Il27ra*^-/-^ mice (scale bar: 300 μm). See Supplemental Figure 5 for quantified data.

To extend the FIMO analysis of STAT transcription factor sites, we used MEME Suite to identify promoter motifs for STAT1, STAT2, STAT3, STAT4, STAT5 (comprising STAT5a and STAT5b), and STAT6. Analysis showed that *Il6ra* deficiency restricted chromatin access to all STAT transcription factors at day 10 of AIA (**Figure 8B**). While promoter regions for each STAT transcription factor remained accessible across the entire course of AIA in WT and *Il27ra*^-/-^ mice, we noted some alterations in the frequency of sites for STAT transcription factors that share a regulatory interplay or cross-regulation (e.g., STAT1 and STAT3, and STAT5 and STAT6) (**Figure 8B** & **8C**) (*25, 27, 31, 38, 41, 44, 45*). For example, WT and *Il27ra*^-/-^ mice with AIA showed similarities in the availability of STAT1 and STAT3 consensus binding sequences at days 3 and 10 of synovitis. However, access to STAT1 sites appeared more transient, with their frequency reduced by day 10 (**Figure 8B**). This alteration in the balance of available STAT1 and STAT3 sites may impact the regulation of genes controlling synovitis in AIA.

To test the link between STAT1 and STAT3 in synovitis, WT and *Il27ra*^-/-^ mice were treated (i.a.) with CpG-*Stat3*-siRNA at the initiation of AIA (**Figure 8D**). This intervention modality targets TLR9-positive cells, delivering a siRNA that silences *Stat3* expression (*42, 43*). Immunohistochemistry showed that TLR9-positive cells populate the synovial lining and sublining of the inflamed synovium from mice with AIA (Supplemental Figure 5). Analysis of synovial mRNA by quantitative PCR confirmed that CpG-*Stat3*-siRNA dampened the synovial expression of *Stat3* (Supplemental Figure 5). CpG-*Stat3*-siRNA treatment improved histological scores for synovial infiltration, synovial hyperplasia, and cartilage and bone erosion in AIA (**Figure 8D**). RNA-seq of synovial tissue from WT and *Il27ra*^-/-^ mice showed that CpG-*Stat3*-siRNA altered synovial gene expression at days 3 and 10 of AIA (**Figure 8E**; Supplemental Figure 5). These include the suppression of genes affecting the deposition of extracellular matrix (e.g., *Adamts17*, *Col9a1*, *Col9a2, Col10a1*), bone and cartilage turnover (e.g., *Dlk1*, *Mdk*, *Gdf5*), and fibroblast activation (e.g., *Fgfrl1*, *Fgfr2*, *Tbx2*, *Ngfr*) (**Figure 8E**). Illustrating the importance of STAT3 in guiding synovial heterogeneity, we identified that CpG-*Stat3*-siRNA treatment of *Il27ra*^-/-^ mice reduced the expression of transcripts involved in the development, maintenance, and activity of ELS (e.g., *Cxcl13*, *Ccl19*, *Ltb*, *Vegfc*) (**Figure 8F**). Consistent with this, immunohistochemistry of the inflamed synovium of *Il27ra*^-/-^ mice showed a reduced incidence of ELS following CpG-*Stat3*-siRNA treatment (**Figure 8G**; Supplemental Figure 5).

To test whether STAT1 and STAT3 share a regulatory interplay in synovitis, we performed a gene set enrichment analysis (GSEA) of transcripts expressed during AIA following CpG-*Stat3*-siRNA treatment. GSEA of RNA-seq data from WT and *Il27ra*^-/-^ mice treated with CpG-*Stat3*-siRNA showed evidence of STAT1/STAT3 cross-regulation, with the inhibition of STAT3-regulated genes associated with an increase in genes controlled by STAT1 (**Figure 9A**). Evidence for STAT1/STAT3 cross-regulation was also obtained with upstream regulator analysis in IPA (**Figure 9B**). A heatmap of this analysis provides examples of genes targeted by CpG-*Stat3*-siRNA in WT and *Il27ra*^-/-^ mice with AIA (**Figure 9C**).

**Figure 9.**
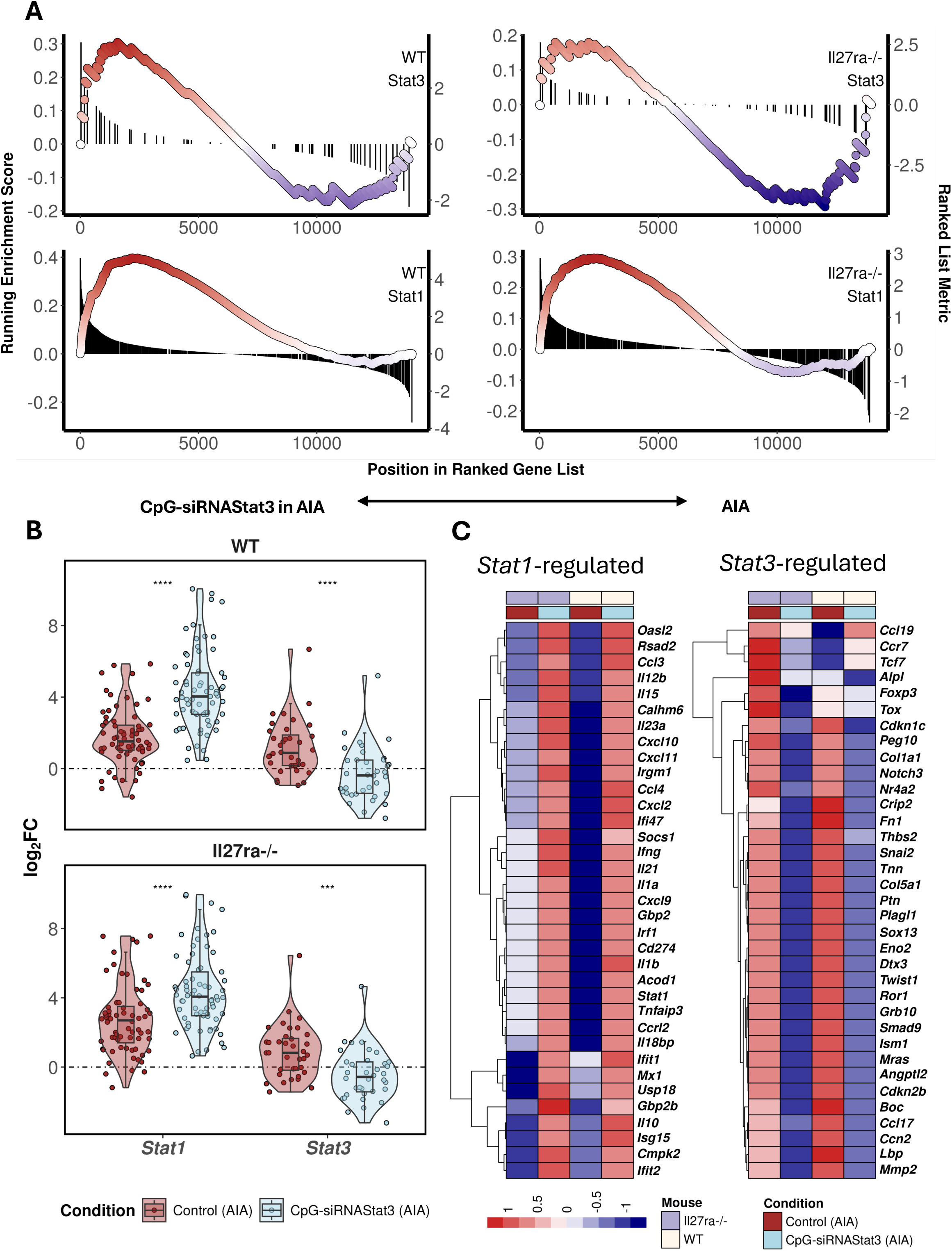
CpG-*Stat3*-siRNA enhances STAT1 signaling in WT and *Il27ra*^-/-^ mice with AIA. **(A)** GSEA of RNA-seq data from the synovium of WT and *Il27ra*^-/-^ mice with AIA. Data show the response to CpG-*Stat3*-siRNA treatment for transcripts affiliated with STAT1 or STAT3. Gene sets used for GSEA were extracted using DoRothEA (decoupleR). **(B)** Violin plots show relative changes in synovial *Stat1* or *Stat3* expression as a response to CpG-*Stat3*-siRNA in AIA at days 3 and 10 of AIA. **(C)** Heatmap showing examples of genes differentially regulated by CpG-*Stat3*-siRNA (Log2FC>1; adjPvalue<0.05).

Reflecting on the increase in STAT1 target genes as a response to CpG-*Stat3*-siRNA treatment, we examined their relationship to leukocyte activation markers. The circos plots show that CpG-*Stat3*-siRNA treatment of WT and *Il27ra*^-/-^ mice altered the proportion of STAT1 target genes in lymphocytes and monocytic cells (**Figure 10A**). Heatmaps provide examples of genes affected by CpG-*Stat3*-siRNA treatment (**Figure 10A**). Focusing on the control of AIA by CpG-*Stat3*-siRNA in *Il27ra*^-/-^ mice (**Figure 9C** & **10A**), we observed patterns of gene regulation linked to lymphocyte function. These included cytokines (e.g., *Ifng*, *Il1b, Il10, Il12b, Il15*), signaling intermediates (e.g., *Stat1*, *Irf1*, *Socs1*), interferon-inducible chemokines (e.g., *Cxcl9*, *Cxcl10*, *Cxcl11*), interferon-stimulated genes (e.g., *Isg15*, *Gbp2*, *Mx1*, *Oasl2*), and negative regulators of innate immunity (e.g., *Il18bp*, *Tnfaip3*). Conversely, our analysis showed that CpG-*Stat3*-siRNA suppresses genes associated with ELS formation (e.g., *Angptl2*, *Ccl17*, *Ccl19, Ccr7*) (**Figure 9C** & **10A**). In this regard, immunohistochemistry of the inflamed synovium of *Il27ra*^-/-^ mice with AIA showed that CpG-*Stat3*-siRNA disrupted the development of synovial ELS (Supplemental Figure 5). Thus, synovial STAT1 and STAT3 share a regulatory interplay during synovitis. To illustrate this point, ranked enrichment plots show the impact of CpG-*Stat3*-siRNA on pathways affecting synovitis in WT and *Il27ra*^-/-^ mice with AIA (**Figure 10B**).

**Figure 10.**
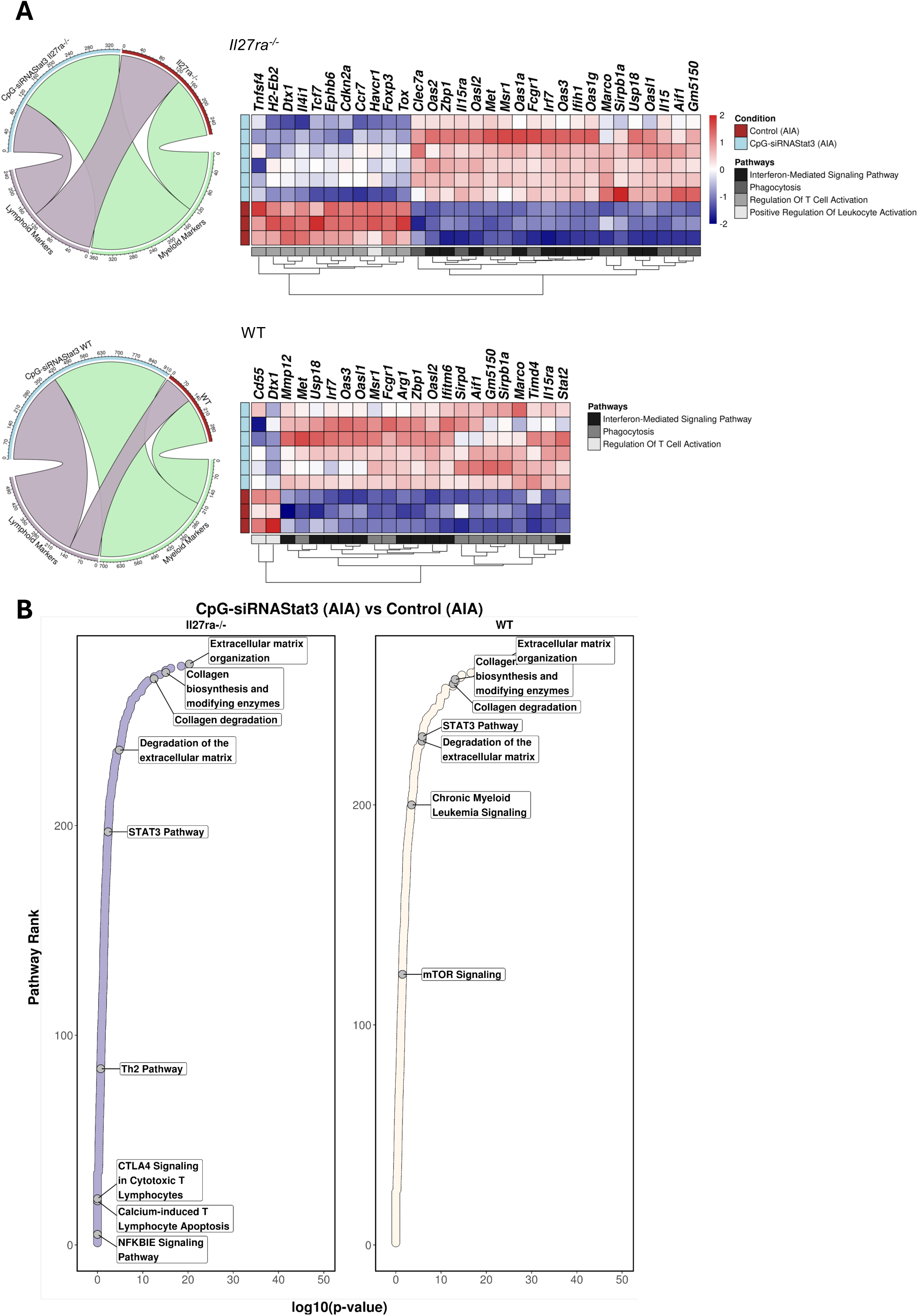
CpG-*Stat3*-siRNA impacts defined signaling mechanisms in WT and *Il27ra*^-/-^ mice with AIA. **(A)** Circos plot maps synovial transcripts differentially regulated by CpG-*Stat3*-siRNA against gene lists characteristic of myeloid or lymphoid cells. Data is shown for WT and *Il27ra*^-/-^ mice with AIA. Genes extracted from the circos plot are presented as a heatmap depicting changes in the expression of myeloid and lymphoid phenotypic markers and associated pathways (clusterprofileR). **(B)** Ranked enrichment plots identify molecular pathways (IPA) inhibited by CpG-*Stat3*-siRNA in WT and *Il27ra*^-/-^ mice with AIA (at day 10 of disease).

### STAT1 and STAT3 signaling provide insight into synovitis heterogeneity in rheumatoid arthritis

To investigate whether an interplay between STAT1 and STAT3 influences the transcriptional programs of synovitis in humans, we first examined the expression of STAT transcription factors in RNA-seq datasets from joint biopsies categorized as fibroblast-rich, myeloid-rich, or lymphoid-rich. Focusing on the roles of STAT1 and STAT3, we correlated their synovial expression levels with SOCS1 and SOCS3 as surrogate markers of their activity (**Figure 11A**). While STAT3 expression was similar across all three synovial pathotypes, a more distinct pattern emerged for STAT1, with the lowest expression in fibroblast-rich synovitis and the highest in lymphoid-rich synovitis (**Figure 11A**). SOCS1 and SOCS3 act as negative feedback inhibitors of STAT1 and STAT3 activation, respectively, and as STAT-inducible factors, they provide insights into the activation states of these transcription factors. Synovial SOCS1 strongly correlated with STAT1 in lymphoid-rich and fibroblast-rich synovitis (**Figure 11A**), though this relationship was less prominent in myeloid-rich synovitis. Conversely, myeloid-rich synovitis showed a closer association with STAT3 activity, as evidenced by a positive correlation between synovial SOCS3 and STAT3 (**Figure 11A**). These results indicate a potential imbalance in STAT1 and STAT3 signaling across different synovial pathotypes. To illustrate this, we performed GSEA on RNA-seq data from patients with myeloid-rich and lymphoid-rich synovitis to assess target gene regulation for STAT1 and STAT3 (**Figure 11B**). While STAT3 target gene expression remained relatively consistent between the two pathotypes, STAT1 activity signatures were more prominent in lymphoid-rich synovitis (**Figure 11B**). Therefore, alterations to the balance of STAT1 and STAT3 signaling within the inflamed joint may impact transcriptional programs, influencing the heterogeneity of synovitis in rheumatoid arthritis.

**Figure 11.**
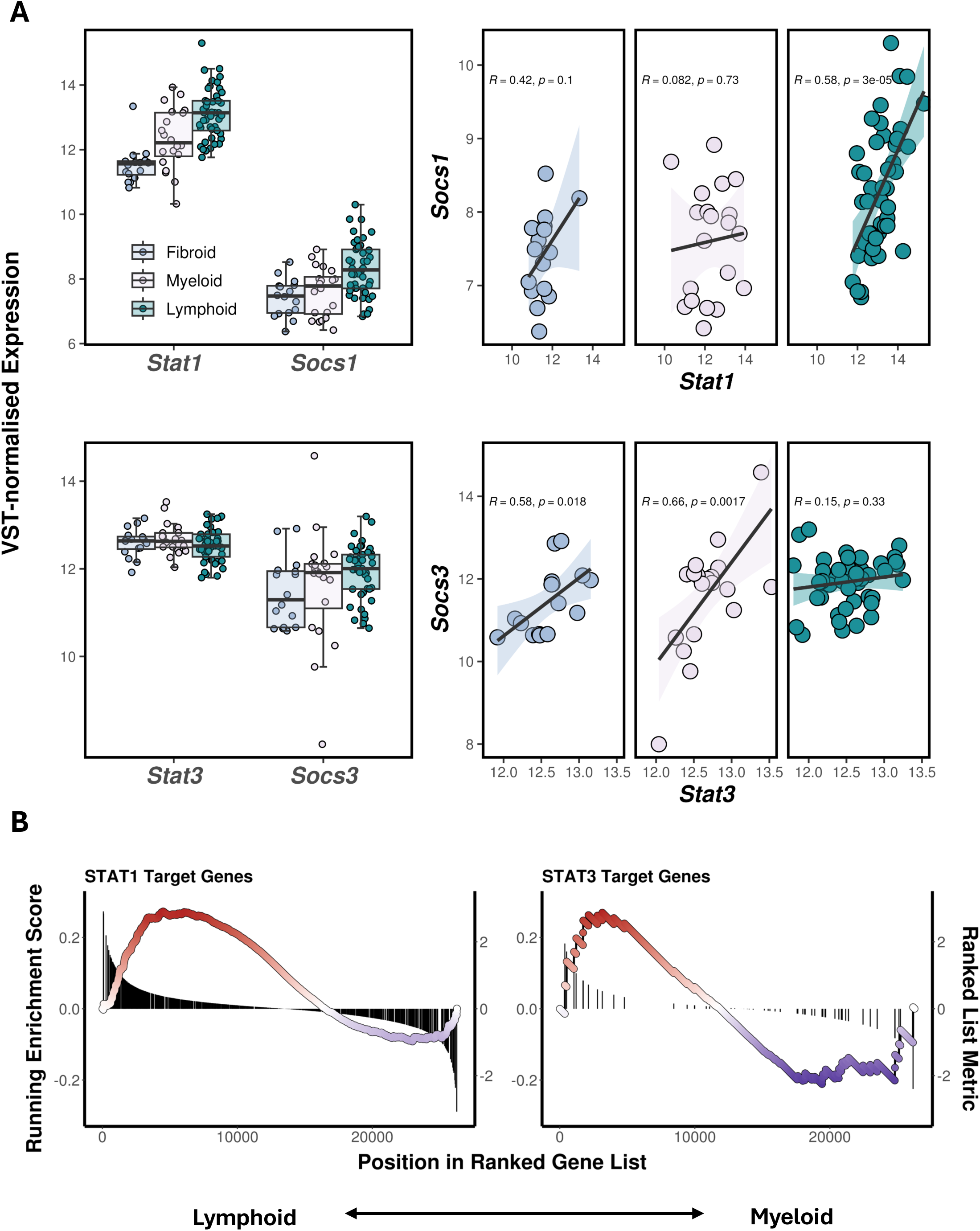
CpG-*Stat3*-siRNA impacts defined signaling mechanisms in WT and *Il27ra*^-/-^ mice with AIA. **(A)** Synovial tissue acquired through a minimally invasive ultrasound synovial biopsy from early arthritis patients was analyzed by RNA-seq (E-MTAB-6141). Synovial expression of *STAT1* and *SOCS1* (top panels) and *STAT3* and *SOCS3* (bottom panels) is shown for tissues with fibroblast-rich (FR; n=17), myeloid-rich (MR; n=21), or lymphoid-rich synovitis (LR; n=46). Correlation plots illustrate the relationship between *STAT1* and *SOCS1* and *STAT3* and *SOCS3* expression in each synovial pathotype. Statistics (Pearson’s r^2^) are given for each. **(B)** GSEA of STAT1 and STAT3 target genes identified using DoRothEA (decoupleR) in synovial RNA-seq from patients with rheumatoid arthritis (E-MTAB-6141). Analysis focused on comparisons between tissues with myeloid-rich and lymphoid-rich synovitis.

## Discussion

Rheumatoid arthritis remains a complex and challenging disease to treat, with patients commonly showing inadequate responses to certain drug classes (*1–3*). These include medicines that target cytokine activities controlling inflammatory joint disease, autoimmunity, comorbidities, and patient multimorbidity (*1, 2, 46, 47*). Thus, there is a need to identify the contributing pathways that influence the progression of arthritis in patients whose disease is undifferentiated at presentation.

Synovial biopsies display histological features that broadly classify pathotypes as fibroblast-rich (pauci-immune), myeloid-rich, or lymphoid-rich synovitis (*9, 11, 13, 20, 48, 49*). These affect responses to biological and targeted therapies. Patients with lymphoid-rich synovitis often exhibit severe disease and poorly respond to anti-TNF therapies (*50*). This has led to clinical trials assessing the effectiveness of biological medicines in patients stratified according to synovial pathotype (*4, 6, 9, 10, 49*). However, several questions remain. These questions concern how synovitis develops and whether synovial pathotypes originate from a common mechanism or unique inflammatory cues. Studies of synovial pathotypes report that the histological features of synovitis reflect the transcriptional hallmarks of synovial gene expression (*5, 7, 9*). These include markers of invasive myofibroblasts and activated synoviocytes in fibroblast-rich synovitis, or lymphoneogenesis in lymphoid-rich synovitis (*5, 7, 9, 14, 16*). However, the regulatory mechanisms directing these disease processes are unclear. Combining data from synovial biopsies from patients with rheumatoid arthritis and mice with AIA, we now provide new insights into the control of synovitis heterogeneity.

The Jak-STAT pathway is a critical determinant in synovitis and is a target for cytokine inhibitors used in rheumatoid arthritis (*1, 2*). Studies show that Jak-STAT signaling orchestrates leukocyte infiltration, controlling their effector characteristics and maintenance within the inflamed synovium (*17, 34, 35, 51*). This pathway also regulates osteoclastogenesis, cartilage erosion, and synovial hyperplasia (*17, 52, 53*). Our data from *Il6ra*^-/-^ and *Il27ra*^-/-^ mice with AIA reaffirm these findings and highlight roles for STAT1 and STAT3 transcription factors in shaping synovitis (*54*). Compared to IL-6, IL-27 is a more potent activator of STAT1, often inhibiting STAT3-driven responses to limit T-cell-mediated pathology and the programming of monocytic cells (*55–58*). In inflammatory models of arthritis, IL-27 inhibits osteoclastogenesis and associated joint pathology, with *Il27ra*^-/-^ mice displaying a more severe form of synovitis than WT mice (*59–61*). This phenotype contrasts with the low inflammatory synovitis evident in *Il6ra*^-/-^ mice and, as previously noted, in *Il6*^-/-^ mice (*19, 32, 34, 36, 37*). Building on these findings, we now show that repeated flares of AIA in *Il6*^-/-^ mice promote synovial hyperplasia, despite lacking an immune cell infiltrate. Thus, two cytokines with shared but contrasting biology develop two distinct forms of synovitis in AIA.

The transcriptional properties of STAT1 and STAT3 are often linked, with STAT1 influencing or suppressing STAT3-mediated gene regulation (*25, 34, 38, 41, 44, 45, 58*). This transcriptional interplay (cross-regulation) controls T-cell effector responses and a shift from competent host defense to inflammation-induced tissue injury (*25, 38, 44*). In experimental models of arthritis, STAT1 plays a protective role, with STAT3 activity promoting synovial infiltration, pannus formation, and joint damage (*34, 35, 41, 51, 62*). Our findings suggest that the balance of STAT1 and STAT3 signaling in the inflamed joint may be critical to the development of myeloid-rich and lymphoid-rich synovitis, with the inhibitory action of CpG-*Stat3*siRNA offering evidence of STAT1/STAT3 cross-regulation. This feature of IL-6 biology has been observed in other inflammatory settings (*44, 63*). However, several questions remain. *How is the balance between STAT1 and STAT3 signaling regulated*? Potential mechanisms include changes to the cytokine network with the inflammatory milieu, or intracellular regulatory mechanisms acting as rheostats of Jak-STAT signaling (*24, 38, 44*). Modulators of cytokine signaling include protein tyrosine phosphatases (e.g., PTPN2, PTPN11, PTPN22, DUSP2) and signaling differences elicited by individual Jak proteins (*38, 64–67*). PTPN2 is highly expressed in lymphoid-rich synovitis, with PTPN2 inhibiting STAT1 signaling to modify the transcriptional output of activated naïve and effector memory CD4^+^ T-cells (*38*). These include genes showing heightened synovial expression in *Il27ra*^-/-^ mice with AIA, such as *Il17a*, *Il17f*, *Il21*, *Cd274,* and homeostatic chemokines and chemokine receptors (*20*). *What is the impact of STAT1 on synovial pathology when STAT3 is inhibited*? Studies involving the genetic ablation of *Stat1* or *Stat3*, or methods that manipulate the expression of SOCS1 and SOCS3, support the primary actions of STAT1 and STAT3 transcription factors in synovitis (*34, 51, 68, 69*). The reality is probably more nuanced, with evidence suggesting that STAT1 becomes more prominent in the latter, more chronic stages of disease (*34, 51*). Our previous studies of *gp130*^Y757F:Y757F^ mice with AIA illustrate this complexity (*35, 41*). These mice possess a single tyrosine-to-phenylalanine substitution in the cytoplasmic domain of gp130, causing a prolonged activation of STAT1 and STAT3 following stimulation by IL-6 family cytokines (including IL-6 and IL-27) (*70–72*). Compared to wild-type littermates, *gp130*^Y757F:Y757F^ mice develop a severe synovitis defined by a more sustained inflammatory infiltrate and worsened joint pathology (*35, 41*). A genetic ablation of *Stat1* (*gp130*^Y757F:Y757F^:*Stat1*^-/-^ mice) or *Stat3* (*gp130*^Y757F:Y757F^:*Stat3*^+/-^ mice) dramatically improved joint pathology in both compound mice (*35, 41*). However, beyond the joint, *Stat1* deletion promoted an increase in effector T-cells, commonly associated with STAT3-driven pathology (*35*). Further research is, therefore, required to understand the extra-articular relationship with STAT1 and STAT3, the consequence of these interactions on synovial pathology, and the role of STAT1 in determining longer-term chronicity and joint pathology. *What drives the pathogenesis of pauci-immune/low inflammatory synovitis in Il6ra^-/-^ mice with AIA*? During synovitis, IL-6 signaling relies on IL-6 trans-signaling, since structural cells within the joint lack IL-6R expression (*19, 73*). This process requires sIL-6R, with synovial sIL-6R levels correlating with leukocyte infiltration and joint damage (*19, 73, 74*). From our data, it is tempting to speculate that the reduced bioavailability of synovial sIL-6R may impact the control of epigenetic mechanisms within the stromal compartment. *Il6ra^-/-^* mice with AIA showed evidence of genome modifications affecting chromatin accessibility to STAT transcription factors and the transcriptional repressor REST in ATAC-seq datasets. Studies from cancer and neuroendocrinology report that REST directly represses STAT1 signaling, with IL-6 accelerating the degradation of REST *via* the ubiquitin-proteasome pathway (*75, 76*). So, what cytokine mechanisms support the development of fibroblast-rich (pauci-immune) synovitis? Our analysis of human biopsies and tissues from *Il6ra*^-/-^ mice with AIA showed enhanced synovial expression of IL-11, IL-13, and Tyk2, commonly associated with fibroblast-driven or fibrotic diseases (*77, 78*). We also saw evidence of regulatory pathways involving signaling intermediates (e.g., WNT, DKK, TR/RXR, AMPK) controlling tissue homeostasis and the activation of fibroblasts, chondrocytes, osteoblasts, and osteoclasts (*77–80*).

Studies of the cellular and molecular mechanisms responsible for synovitis heterogeneity are currently limited by a lack of animal models that generate new hypotheses for testing in humans. The clinical evaluation of biological medicines has begun to illustrate the benefit of stratifying patient treatment according to synovial pathology (*4, 6, 7, 10*). However, further studies are required to extend research beyond correlations. For example, data obtained from human joint biopsies reveal an inverse relationship between synovial *IL27* (p28 subunit) expression and lymphoid-rich synovitis (*20*). The occurrence of synovial ELS in the inflamed joints of *Il27ra*^-/-^ mice with AIA supports this observation and provides a rationale for linking human and mouse datasets. Appreciating the limitations of mouse models in the study of rheumatoid arthritis, studies of *Il6ra*^-/-^ and *Il27ra*^-/-^ mice with AIA provide opportunities to explore how interventions (as exemplified by CpG-*Stat3*siRNA) may modify the course of synovitis in humans. Thus, the model systems described herein open new avenues into the mechanisms controlling the cellular and molecular hallmarks of human synovial biopsies, allowing the generation of new hypotheses that support treatment decisions in patients with inflammatory arthritis.

## Supporting information

Supplemental data and statistical summaries

## Acknowledgements

Research was supported by grants from UKRI-MRC (Reference MR/X00077X/1), Versus Arthritis (References 20770, 19796, 20305, 22706), Wellcome Trust (Reference 107964/Z/15/Z), and the National Health and Medical Research Council (NHMRC) of Australia. BJJ holds a Senior Research Fellowship from the NHMRC. STOH and BCC received PhD Studentships through donations to The Systems Immunity Research Institute from Sir Stanley Thomas. DGH received a PhD Studentship from the GW4 BioMed UKRI-MRC Doctoral Training Partnership. Versus Arthritis, the School of Medicine, and the College of Biomedical Life Sciences at Cardiff University provided joint funding for a PhD Studentship for ADS. The authors thank Drs. David Millrine, Kevin Brennan, and Seva Makeev for bioinformatic advice.

## Supplemental Figures

**Supplemental Figure 1. Disease heterogeneity in synovitis and additional indices of inflammatory arthritis in AIA. (A)** Histological features of human synovial biopsies characterized by immunohistochemistry using antibodies defining B-cells (anti-CD20), T-cells (anti-CD3), monocytic cells (anti-CD68), and plasma cells (anti-CD138). **(B)** Immunohistochemistry of CD3 in synovial tissues from mice with AIA (extracted at day 10 of synovitis). **(C)** Measurement of mBSA antibody titers in serum samples from mice with AIA. **(D)** Changes in joint swelling as a response to AIA onset. Values (mean ± SEM; n=4-6 mice/group) are shown across the time course of the disease and reflect differences between the non-inflamed contralateral joint and the joint with AIA. Histological AIA scores illustrating disease induction above representative contralateral joints from WT mice with AIA. **(E)** Mice were challenged with a repeat relapsing model of AIA. Mice were antigen primed with mBSA, with disease onset triggered by administration of mBSA (i.a.) on Days 0, 14, and 28. Histological scores of synovial hyperplasia (Left) and inflammatory cells (Right) are shown from WT and *Il6*^-/-^ mice. Values show scores from three independent assessors blinded to the study design (mean ± SEM; n=4-6 mice/group).

**Supplemental Figure 2. Additional insights from RNA-seq of synovial pathotypes in mice with AIA. (A)** Bulk RNA-seq of synovial tissues from WT, *Il6ra*^-/-^, and *Il27ra*^-/-^ mice at baseline and days 3 and 10 of AIA. Summary details support interpretations presented in Figure 2. Heatmap showing complete datasets for all individual mice included in the analysis, with differentially regulated genes in response to AIA onset presented for days 3 and 10 of synovitis (Log2FC>1; adjPvalue<0.05). Complete linkage created hierarchical gene clustering. **(B)** Relative expression of chemokine ligands and their receptors in RNA-seq data from WT, *Il6ra*^-/-^, and *Il27ra*^-/-^ mice with AIA. **(C)** Heatmaps showing the relative expression of transcriptional regulators and signaling intermediates associated with common sensing pathways in synovial tissues for WT, *Il6ra*^-/-^, and *Il27ra*^-/-^ mice with AIA at days 3 and 10 of AIA. **(D)** Cluster comparison of fibroblast, myeloid and lymphocyte gene sets in RNA-seq from WT, *Il6ra*^-/-^, and *Il27ra*^-/-^ mice with AIA (from days 3 and 10). Analysis was restricted to transcripts showing altered expression relative to naïve non-challenged control mice. **(E)** Illustrative examples of signaling events identified by molecular pathway (clusterProfiler) and the relative expression of genes within these cascades in WT, *Il6ra*^-/-^, and *Il27ra*^-/-^ mice with AIA. **(F)** Relative expression of cytokines and their receptors in RNA-seq data from WT, *Il6ra*^-/-^, and *Il27ra*^-/-^ mice with AIA. Data for days 3 and 10 of synovitis are shown. **(G)** Statistical summary of intermediates identified by IPA upstream regulator analysis of RNA-seq data.

**Supplemental Figure 3. Alignment of RNA-seq and ATAC-seq datasets. (A)** Consensus (≥2) ATAC-seq peaks were mapped against the mm10 genome and corresponding RNA-seq from WT, *Il6ra*^-/-^, and *Il27ra*^-/-^ mice with AIA. Alignment compares chromatin accessibility with differential gene regulation for each condition. The ChIPseeker package was used to map differentially regulated genes against the mm10 genome. **(B)** Putative transcription factor motifs associated with sequencing peaks from ATAC-seq datasets were identified using motif enrichment analysis in MEME-Suite. Data support the analysis in Figure 4.

**Supplemental Figure 4. Indices of synovitis following treatment with CpG-*Stat3*-siRNA. (A)** Treatment controls for mice with AIA. At the point of arthritis induction, mice were administered (i.a.) with mBSA alone or in combination with either CpG or non-hybridized single-stranded CpG*-Stat3*-siRNA (ssCpG-*Stat3*-siRNA). Histological scores from two independent assessors are shown (mean ± SEM, n=4-5 mice/group). **(B)** WT and *Il27ra*^-/-^ mice were challenged with mBSA. Following antigen priming, arthritis was induced by administration (i.a) of mBSA alone or with CpG-Stat3-siRNA (0.125 nmol/μl). Total RNA was extracted from the synovium on days 3 and 10 of AIA. *Stat3* expression was monitored by quantitative PCR. **(C)** Immunohistochemistry of CD3 and TLR9 in the inflamed synovium of mice with AIA (scale bar: 500 μm [left]; 200 μm [right]). Digital imaging quantified staining within the field of view (*P*<0.05; n=7 mice/group; mean ± SEM). **(D)** An extended heatmap showing differentially regulated genes in response to CpG-Stat3-siRNA treatment of mice with AIA. **(E)** Image quantification of immunohistochemistry in Figure 5F. The number of synovial lymphocyte aggregates was quantified (*P*<0.05; n = 5-10 sections/group; mean ± SEM).

**Supplemental Figure 5. Comparison of cellular and molecular hallmarks from mouse and human synovial pathotypes. (A)** Overlapping gene sets were analyzed with IPA. The relative expression of genes linked with lymphocyte activation and fibroblast involvement is shown for RNA-seq data from mice with AIA and human synovial samples. **(B)** The hclust function created a correlation matrix between mouse and human RNA-seq data for the genes identified with IPA (A) at days 3 and 10 of AIA.

## Supplemental Tables

**Supplemental Table 1** A Wilcoxon rank sum test was used to conduct pair-wise comparisons between each time point of AIA (day 3 and 10) for each motif, per mouse strain.

**Supplemental Table 2** The Wilcoxon rank sum test was used to conduct pair-wise comparisons between each mouse strain for each motif, restricted to day 3 of AIA.

**Supplemental Table 3** The Wilcoxon rank sum test was used to conduct pair-wise comparisons between each mouse strain for each motif, restricted to day 10 of AIA.

