## Supplemental data and statistical summaries for "Discrete cytokine signaling networks instruct distinct synovial pathotypes in inflammatory arthritis"

#### Supplemental Figure 1

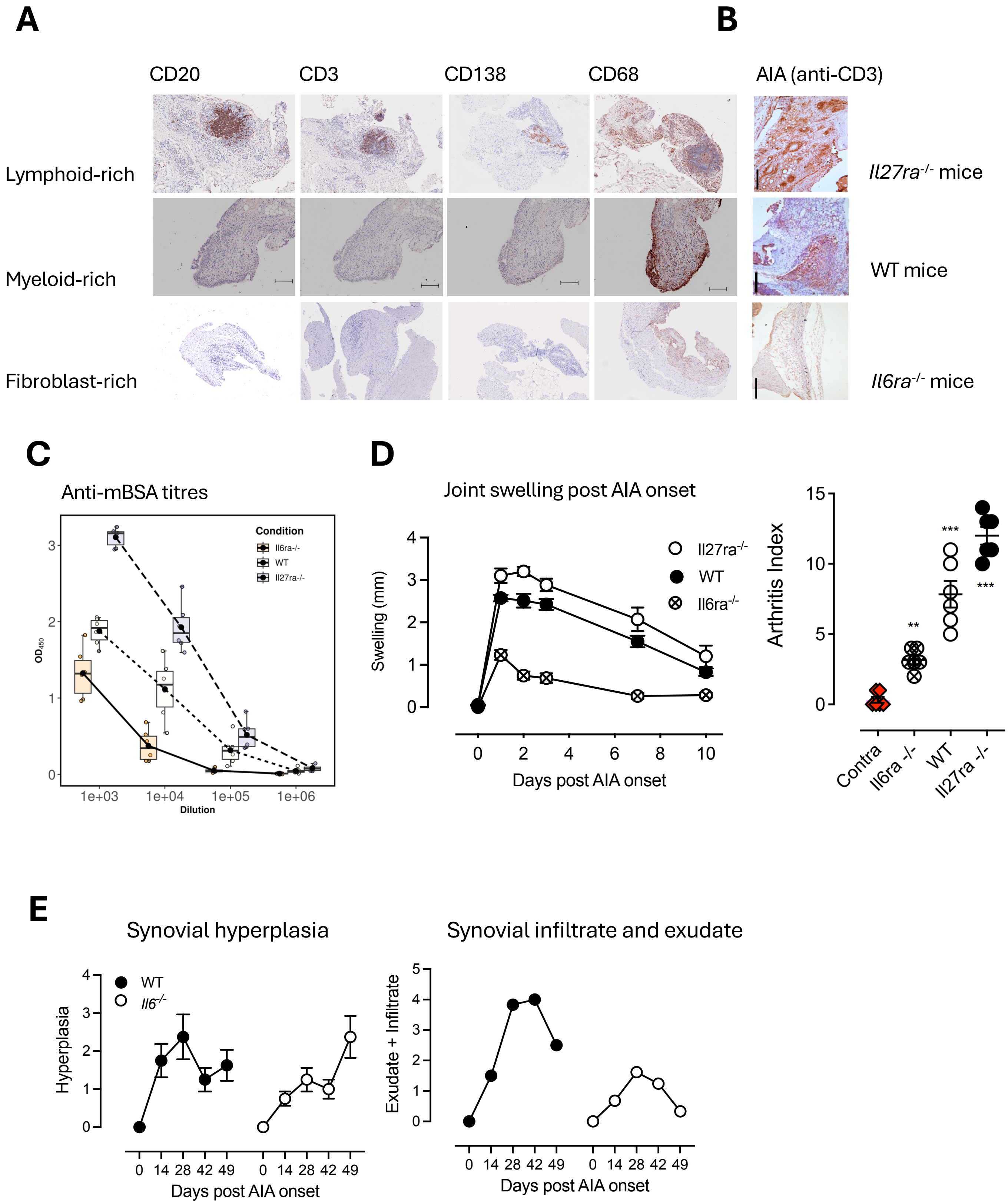

### Supplemental Figure 2

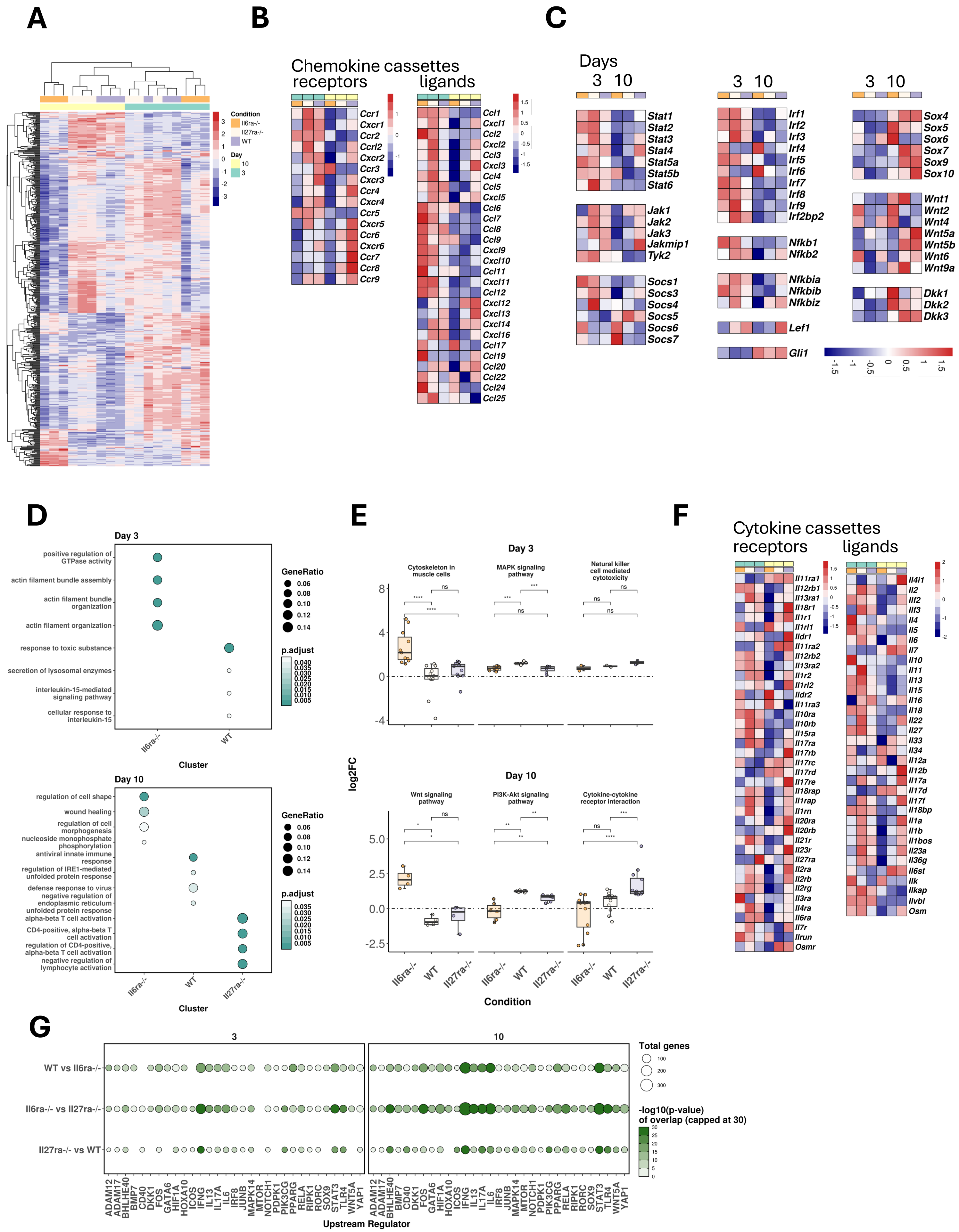

#### Supplemental Figure 3

*Il6ra-/-* *WT* *Il27ra-/-*

***Day 3***

***Day 10***

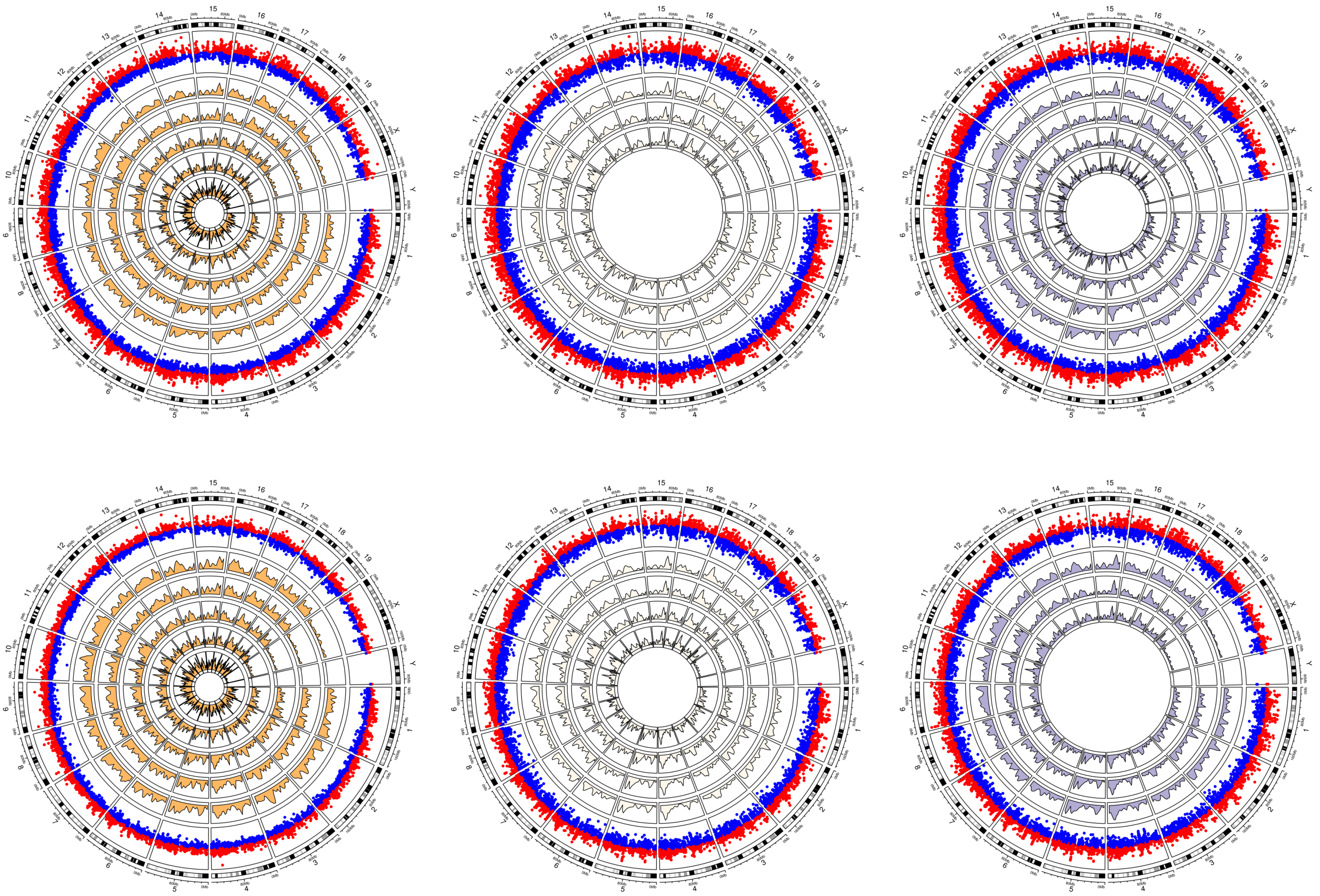

**AP-1**

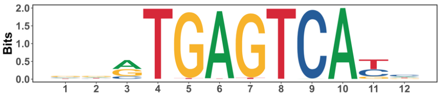

**BACH**

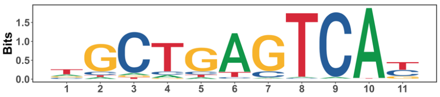

**CREB**

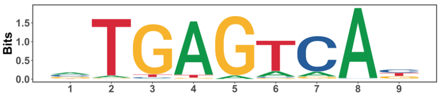

**CREB**

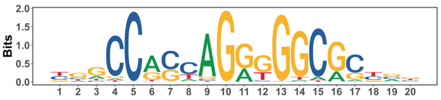

**CTCF**

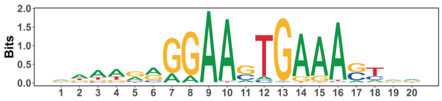

**IRF**

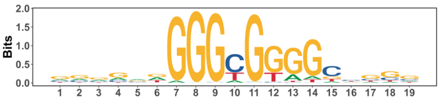

**KLF**

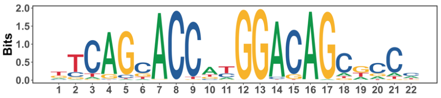

**REST**

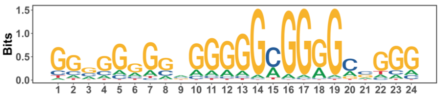

**SP**

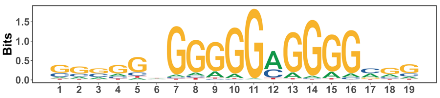

**ZBTB**

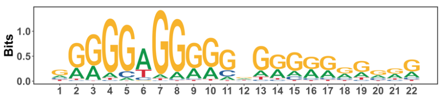

**E2F**

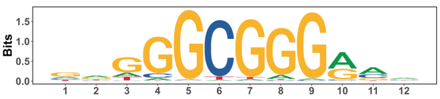

**ETS**

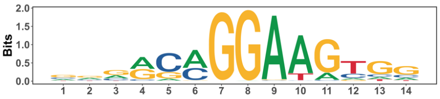

### Supplemental Figure 4

A

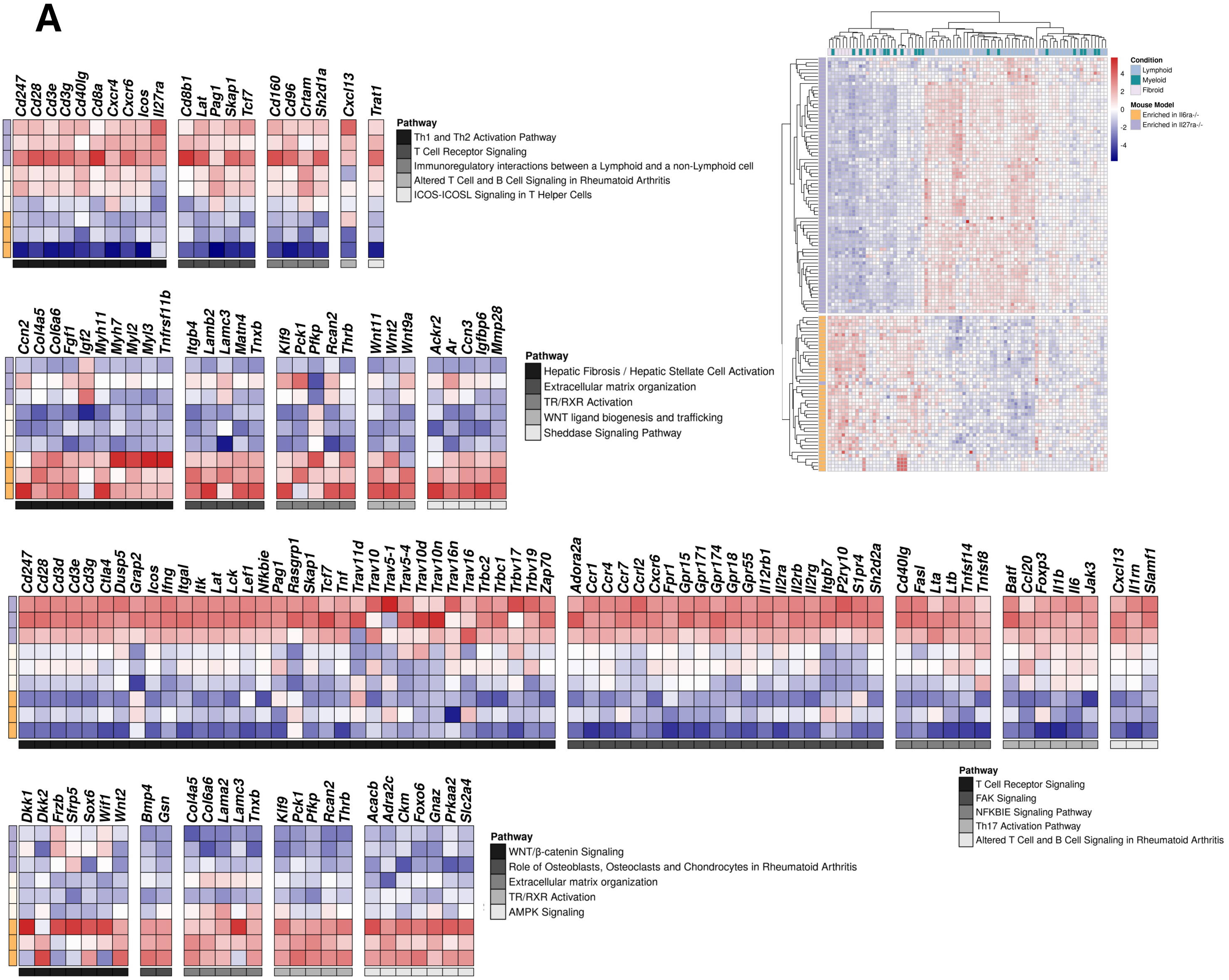


B

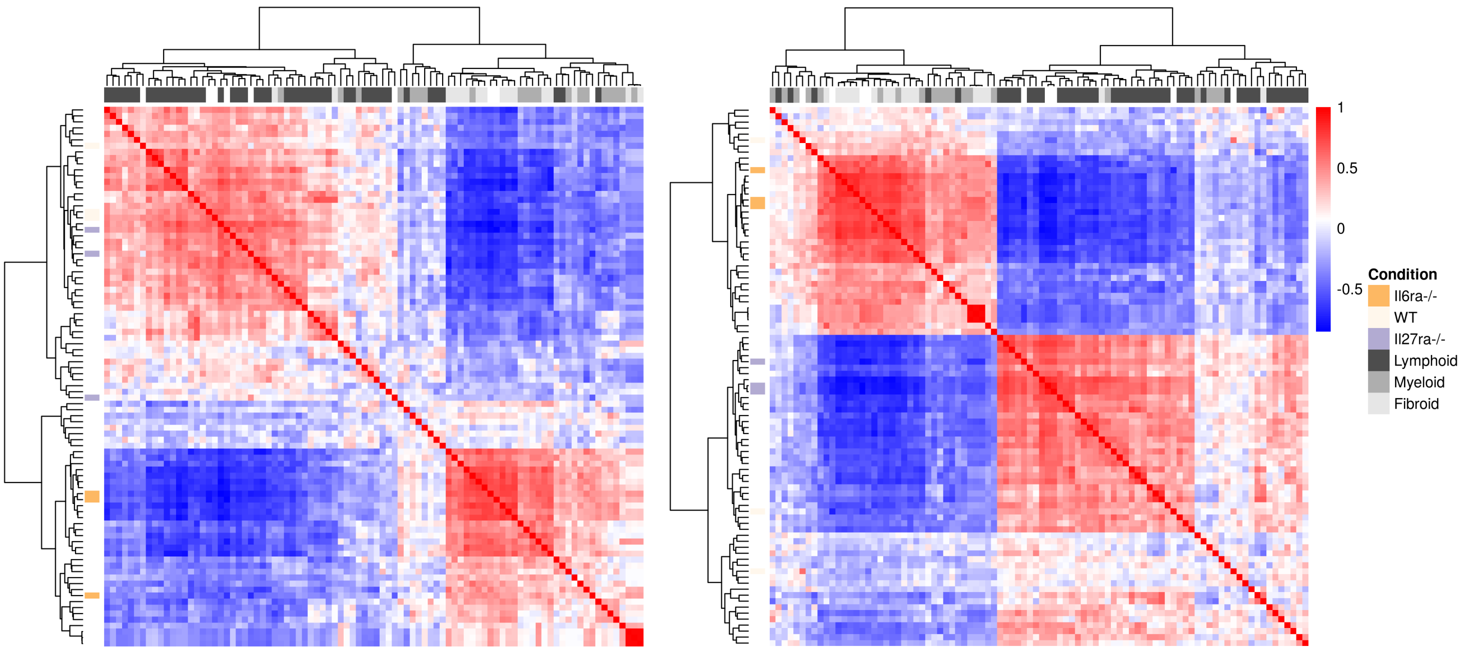

#### Supplemental Figure 5

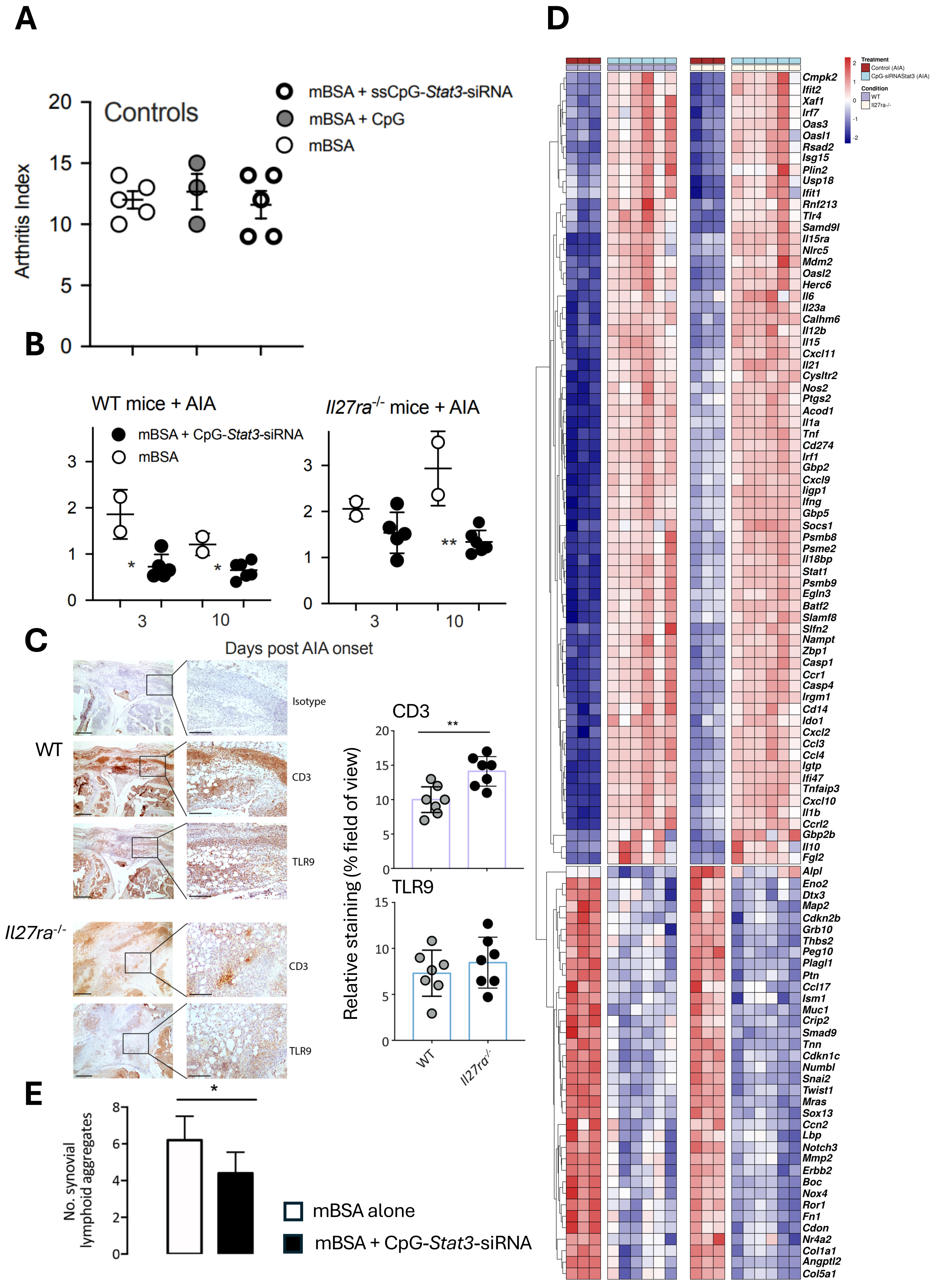

Motif Enrichment Results (Day 3)

| TF Motif | Comparison | W | p-value | adjusted p-value |
|---|---|---|---|---|
| STAT1 | Il6ra-/- vs WT | 3087948.0 | 1.657069e-27 | 4.971207e-27 |
| STAT1 | Il6ra-/- vs Il27ra-/- | 3181352.0 | 3.892320e-01 | 3.892320e-01 |
| STAT1 | WT vs Il27ra-/- | 2515367.5 | 2.161139e-21 | 3.241708e-21 |
| MYOD1 | Il6ra-/- vs WT | 7790566.0 | 7.819967e-304 | 2.345990e-303 |
| MYOD1 | Il6ra-/- vs Il27ra-/- | 7202698.0 | 1.007621e-191 | 1.511432e-191 |
| MYOD1 | WT vs Il27ra-/- | 1989983.5 | 5.235736e-42 | 5.235736e-42 |
| MYOG | Il6ra-/- vs WT | 10792346.5 | 0.000000e+00 | 0.000000e+00 |
| MYOG | Il6ra-/- vs Il27ra-/- | 10104498.5 | 0.000000e+00 | 0.000000e+00 |
| MYOG | WT vs Il27ra-/- | 2252286.0 | 4.282649e-69 | 4.282649e-69 |
| STAT2 | Il6ra-/- vs WT | 2135451.0 | 2.224273e-06 | 2.224273e-06 |
| STAT2 | Il6ra-/- vs Il27ra-/- | 2384584.0 | 2.250568e-85 | 6.751703e-85 |
| STAT2 | WT vs Il27ra-/- | 1494624.5 | 7.919096e-74 | 1.187864e-73 |
| PPARG | Il6ra-/- vs WT | 2686950.0 | 3.112553e-194 | 9.337659e-194 |
| PPARG | Il6ra-/- vs Il27ra-/- | 2567237.5 | 7.361930e-184 | 1.104289e-183 |
| PPARG | WT vs Il27ra-/- | 770946.0 | 1.014535e-06 | 1.014535e-06 |
| SOX3 | Il6ra-/- vs WT | 508590.0 | 1.563092e-02 | 1.563092e-02 |
| SOX3 | Il6ra-/- vs Il27ra-/- | 592542.5 | 1.831191e-09 | 2.746787e-09 |
| SOX3 | WT vs Il27ra-/- | 377347.5 | 5.047475e-16 | 1.514242e-15 |
| GLI1 | Il6ra-/- vs WT | 5786352.0 | 0.000000e+00 | 0.000000e+00 |
| GLI1 | Il6ra-/- vs Il27ra-/- | 5762868.0 | 0.000000e+00 | 0.000000e+00 |
| GLI1 | WT vs Il27ra-/- | 64541.0 | 0.000000e+00 | 0.000000e+00 |
| STAT3 | Il6ra-/- vs WT | 523034.0 | 6.803112e-66 | 1.020467e-65 |
| STAT3 | Il6ra-/- vs Il27ra-/- | 481156.0 | 3.328459e-73 | 9.985376e-73 |
| STAT3 | WT vs Il27ra-/- | 560259.5 | 2.577636e-01 | 2.577636e-01 |
| STAT4 | Il6ra-/- vs WT | 380538.0 | 4.459607e-21 | 8.988192e-21 |
| STAT4 | Il6ra-/- vs Il27ra-/- | 351551.0 | 5.992128e-21 | 8.988192e-21 |
| STAT4 | WT vs Il27ra-/- | 283900.0 | 1.800307e-02 | 1.800307e-02 |
| P63 | Il6ra-/- vs WT | 2838187.0 | 0.000000e+00 | 0.000000e+00 |
| P63 | Il6ra-/- vs Il27ra-/- | 2851678.0 | 0.000000e+00 | 0.000000e+00 |
| P63 | WT vs Il27ra-/- | 618643.0 | 7.379903e-13 | 7.379903e-13 |
| SNAI1 | Il6ra-/- vs WT | 824840.0 | 2.574280e-212 | 7.722840e-212 |
| SNAI1 | Il6ra-/- vs Il27ra-/- | 803006.0 | 2.242484e-206 | 3.363725e-206 |
| SNAI1 | WT vs Il27ra-/- | 884780.0 | 1.022744e-38 | 1.022744e-38 |
| STAT5 | Il6ra-/- vs WT | 567229.5 | 4.993142e-67 | 1.497943e-66 |
| STAT5 | Il6ra-/- vs Il27ra-/- | 531403.0 | 4.671601e-60 | 7.007402e-60 |
| STAT5 | WT vs Il27ra-/- | 628226.0 | 4.647600e-01 | 4.647600e-01 |
| STAT6 | Il6ra-/- vs WT | 326598.0 | 3.256528e-09 | 4.884793e-09 |
| STAT6 | Il6ra-/- vs Il27ra-/- | 349819.0 | 9.882229e-01 | 9.882229e-01 |
| STAT6 | WT vs Il27ra-/- | 286244.0 | 2.083093e-18 | 6.249279e-18 |

Motif Enrichment Results (Day 10)

| TF Motif | Comparison | W | p-value | adjusted p-value |
|---|---|---|---|---|
| STAT1 | Il6ra-/- vs WT | 109504.5 | 7.283611e-286 | 2.185083e-285 |
| STAT1 | Il6ra-/- vs Il27ra-/- | 63963.0 | 4.032422e-204 | 6.048633e-204 |
| STAT1 | WT vs Il27ra-/- | 437393.5 | 1.348554e-01 | 1.348554e-01 |
| MYOD1 | Il6ra-/- vs WT | 10929667.5 | 2.040062e-272 | 3.060093e-272 |
| MYOD1 | Il6ra-/- vs Il27ra-/- | 6521751.5 | 0.000000e+00 | 0.000000e+00 |
| MYOD1 | WT vs Il27ra-/- | 2254483.5 | 4.612998e-26 | 4.612998e-26 |
| MYOG | Il6ra-/- vs WT | 14891244.5 | 0.000000e+00 | 0.000000e+00 |
| MYOG | Il6ra-/- vs Il27ra-/- | 8095680.5 | 0.000000e+00 | 0.000000e+00 |
| MYOG | WT vs Il27ra-/- | 2978799.5 | 5.546907e-74 | 5.546907e-74 |
| STAT2 | Il6ra-/- vs WT | 10617.5 | 1.384603e-248 | 4.153808e-248 |
| STAT2 | Il6ra-/- vs Il27ra-/- | 5935.0 | 5.287675e-183 | 7.931512e-183 |
| STAT2 | WT vs Il27ra-/- | 201346.5 | 4.077916e-01 | 4.077916e-01 |
| PPARG | Il6ra-/- vs WT | 3426766.0 | 0.000000e+00 | 0.000000e+00 |
| PPARG | Il6ra-/- vs Il27ra-/- | 1963979.0 | 1.581572e-305 | 2.372358e-305 |
| PPARG | WT vs Il27ra-/- | 573892.0 | 2.432124e-08 | 2.432124e-08 |
| SOX3 | Il6ra-/- vs WT | 200742.0 | 2.185016e-56 | 6.555049e-56 |
| SOX3 | Il6ra-/- vs Il27ra-/- | 146165.5 | 3.248641e-13 | 3.248641e-13 |
| SOX3 | WT vs Il27ra-/- | 183148.5 | 1.572578e-23 | 2.358868e-23 |
| GLI1 | Il6ra-/- vs WT | 9024858.0 | 0.000000e+00 | 0.000000e+00 |
| GLI1 | Il6ra-/- vs Il27ra-/- | 5181894.0 | 0.000000e+00 | 0.000000e+00 |
| GLI1 | WT vs Il27ra-/- | 369592.0 | 1.974045e-205 | 1.974045e-205 |
| STAT3 | Il6ra-/- vs WT | 47105.5 | 2.318048e-258 | 6.954144e-258 |
| STAT3 | Il6ra-/- vs Il27ra-/- | 25233.5 | 5.480147e-184 | 8.220221e-184 |
| STAT3 | WT vs Il27ra-/- | 236990.0 | 4.580939e-01 | 4.580939e-01 |
| STAT4 | Il6ra-/- vs WT | 28526.0 | 7.130335e-186 | 2.139101e-185 |
| STAT4 | Il6ra-/- vs Il27ra-/- | 16826.0 | 3.076988e-135 | 4.615483e-135 |
| STAT4 | WT vs Il27ra-/- | 147338.5 | 6.712992e-04 | 6.712992e-04 |
| P63 | Il6ra-/- vs WT | 4155197.0 | 0.000000e+00 | 0.000000e+00 |
| P63 | Il6ra-/- vs Il27ra-/- | 2111100.5 | 7.810920e-193 | 7.810920e-193 |
| P63 | WT vs Il27ra-/- | 131734.5 | 9.950797e-216 | 1.492620e-215 |
| SNAI1 | Il6ra-/- vs WT | 3192490.0 | 1.457767e-18 | 2.186650e-18 |
| SNAI1 | Il6ra-/- vs Il27ra-/- | 2075646.0 | 1.600087e-01 | 1.600087e-01 |
| SNAI1 | WT vs Il27ra-/- | 1359098.0 | 4.723125e-61 | 1.416937e-60 |
| STAT5 | Il6ra-/- vs WT | 41916.5 | 9.559937e-220 | 2.867981e-219 |
| STAT5 | Il6ra-/- vs Il27ra-/- | 58695.0 | 1.408113e-109 | 2.112169e-109 |
| STAT5 | WT vs Il27ra-/- | 207827.5 | 2.172161e-21 | 2.172161e-21 |
| STAT6 | Il6ra-/- vs WT | 19877.5 | 6.165908e-162 | 1.849772e-161 |
| STAT6 | Il6ra-/- vs Il27ra-/- | 11499.5 | 5.824495e-111 | 8.736742e-111 |
| STAT6 | WT vs Il27ra-/- | 98811.0 | 3.645071e-15 | 3.645071e-15 |

Motif Enrichment: Day 3 vs Day 10 per Condition

| TF Motif | Condition | W | p-value | adjusted p-value |
|---|---|---|---|---|
| STAT1 | Il6ra-/- | 3956853.0 | 0.000000e+00 | 0.000000e+00 |
| STAT1 | WT | 2444862.5 | 0.000000e+00 | 0.000000e+00 |
| STAT1 | Il27ra-/- | 1167389.0 | 8.706937e-194 | 1.617003e-193 |
| MYOD1 | Il6ra-/- | 6722750.0 | 0.000000e+00 | 0.000000e+00 |
| MYOD1 | WT | 1367289.5 | 4.048950e-242 | 8.772726e-242 |
| MYOD1 | Il27ra-/- | 1115998.5 | 1.839343e-55 | 2.314012e-55 |
| MYOG | Il6ra-/- | 8978850.0 | 0.000000e+00 | 0.000000e+00 |
| MYOG | WT | 1446388.0 | 0.000000e+00 | 0.000000e+00 |
| MYOG | Il27ra-/- | 1334132.0 | 3.572317e-59 | 4.644012e-59 |
| STAT2 | Il6ra-/- | 1907263.5 | 0.000000e+00 | 0.000000e+00 |
| STAT2 | WT | 1282719.0 | 2.902770e-305 | 7.075501e-305 |
| STAT2 | Il27ra-/- | 641327.5 | 1.876773e-203 | 3.659708e-203 |
| PPARG | Il6ra-/- | 2739178.0 | 7.486272e-48 | 8.847412e-48 |
| PPARG | WT | 895759.5 | 7.125850e-01 | 7.125850e-01 |
| PPARG | Il27ra-/- | 614478.5 | 4.637587e-26 | 4.759628e-26 |
| SOX3 | Il6ra-/- | 1104779.5 | 1.983138e-135 | 3.093695e-135 |
| SOX3 | WT | 384991.0 | 2.209960e-33 | 2.394123e-33 |
| SOX3 | Il27ra-/- | 229305.5 | 1.405488e-59 | 1.890139e-59 |
| GLI1 | Il6ra-/- | 0.0 | 0.000000e+00 | 0.000000e+00 |
| GLI1 | WT | 0.0 | 0.000000e+00 | 0.000000e+00 |
| GLI1 | Il27ra-/- | 20309.5 | 0.000000e+00 | 0.000000e+00 |
| STAT3 | Il6ra-/- | 1496831.5 | 0.000000e+00 | 0.000000e+00 |
| STAT3 | WT | 715530.0 | 2.792137e-53 | 3.402917e-53 |
| STAT3 | Il27ra-/- | 369022.5 | 4.428646e-30 | 4.668032e-30 |
| STAT4 | Il6ra-/- | 924750.0 | 5.839746e-282 | 1.339707e-281 |
| STAT4 | WT | 374783.5 | 1.065362e-36 | 1.222032e-36 |
| STAT4 | Il27ra-/- | 242751.0 | 6.820338e-73 | 9.499757e-73 |
| P63 | Il6ra-/- | 303913.0 | 0.000000e+00 | 0.000000e+00 |
| P63 | WT | 121797.5 | 0.000000e+00 | 0.000000e+00 |
| P63 | Il27ra-/- | 9682.0 | 0.000000e+00 | 0.000000e+00 |
| SNAI1 | Il6ra-/- | 0.0 | 0.000000e+00 | 0.000000e+00 |
| SNAI1 | WT | 741748.0 | 4.438400e-153 | 7.525983e-153 |
| SNAI1 | Il27ra-/- | 388800.0 | 5.772648e-113 | 8.658972e-113 |
| STAT5 | Il6ra-/- | 1185443.0 | 0.000000e+00 | 0.000000e+00 |
| STAT5 | WT | 786768.5 | 1.736199e-154 | 3.077807e-154 |
| STAT5 | Il27ra-/- | 387659.0 | 5.048525e-142 | 8.203853e-142 |
| STAT6 | Il6ra-/- | 621921.0 | 3.785358e-222 | 7.769946e-222 |
| STAT6 | WT | 320225.5 | 5.331092e-76 | 7.700467e-76 |
| STAT6 | Il27ra-/- | 126434.0 | 9.880174e-36 | 1.100934e-35 |
